# ATP activation of peritubular cells drives testicular sperm transport

**DOI:** 10.1101/2020.09.15.298299

**Authors:** David Fleck, Lina Kenzler, Nadine Mundt, Martin Strauch, Naofumi Uesaka, Robert Moosmann, Felicitas Bruentgens, Annika Missel, Artur Mayerhofer, Dorit Merhof, Jennifer Spehr, Marc Spehr

**Author notes:** Charité-Universitätsmedizin Berlin, Neuroscience Research Center, D-10117 Berlin, Germany (Felicitas Bruentgens). These authors contributed equally to this work (David Fleck, Lina Kenzler).

## Abstract

Spermatogenesis, the complex developmental process of male germ cell proliferation, differentiation, and maturation, is the basis of male fertility and reproductive fitness. In the seminiferous tubules of the testes, spermatozoa are constantly generated from spermatogonial stem cells through a stereotyped sequence of mitotic and meiotic divisions. The basic physiological principles, however, that control both maturation and luminal transport of the still immotile spermatozoa within the seminiferous tubules remain poorly, if at all, defined. Here, we show that coordinated contractions of smooth muscle-like testicular peritubular cells provide the propulsive force for luminal sperm transport towards the rete testis and epididymis. Using a mouse model for *in vivo* imaging, we describe and quantify spontaneous tubular contractions and show a causal relationship between peritubular Ca^2+^ waves and peristaltic transport. Moreover, we identify P2 receptor-dependent purinergic signaling pathways as physiological triggers of tubular contractions both *in vitro* and *in vivo*. When challenged with extracellular ATP, transport of luminal content inside the seminiferous tubules displays stage-dependent directionality. We thus suggest that paracrine purinergic signaling coordinates peristaltic recurrent contractions of the mouse seminiferous tubules to propel immotile spermatozoa to the rete testis. Consequently, our findings could have substantial pharmaceutical implications for both infertility treatment and / or male contraception.

## Introduction

Spermatogenesis ranks among the most complex, yet least understood developmental processes in postnatal life. Inside the seminiferous tubules, this intricate course of mass cell proliferation and transformation events generates haploid spermatozoa from diploid spermatogonial stem cells. Along the seminiferous epithelium, spermatogenesis has been conceptualized by attribution of sequential cellular ‘stages’, which progress through coordinated cycles (Russell, 1990). Each spermatogenic cycle completes with the release of immotile spermatozoa from the seminiferous epithelium into the lumen of the tubule. Once detached from the Sertoli cells, sperm must be transported to the *rete testis* and epididymis for final maturation. Precisely regulated tubular transport mechanisms are, thus, imperative for reproduction.

While bulk movement of luminal content has been anecdotally reported (Cross, 1958; Setchell et al., 1978; Worley et al., 1985), no quantitative data on sperm transport within the seminiferous tubules is available. Early *in vitro* observations of apparent minute undulating motions of seminiferous tubule segments (Roosen-Runge, 1951; Suvanto and Kormano, 1970) suggested that smooth muscle-like testicular peritubular cells (TPCs) (Clermont, 1958; Ross, 1967) could mediate contractile tubule movements. This concept has gained widespread support from several, mostly indirect, *in vitro* studies (Ailenberg et al., 1990; Filippini et al., 1993; Miyake et al., 1986; Tripiciano et al., 1996). However, quantitative direct (i.e., live cell) measurements of seminiferous tubule contractions are rare and controversial (Ellis et al., 1978; Harris and Nicholson, 1998; Losinno et al., 2012; Worley and Leendertz, 1988). Moreover, mechanistic *in vivo* evidence is lacking. Here, we demonstrate that, by acting on ionotropic and metabotropic P2 receptors, extracellular ATP activates TPC contractions that trigger directional sperm movement within the seminiferous tubules both *in vitro* and *in vivo*.

## Results

### ATP is a potent TPC stimulus

Accumulating data suggests that purinergic signaling constitutes a critical component of testicular paracrine communication (Fleck et al., 2016; Foresta et al., 1995; Gelain et al., 2003; Poletto Chaves et al., 2006; Veitinger et al., 2011a; Walenta et al., 2018), with Sertoli cells acting as a primary source of ATP secretion (Gelain et al., 2005). Therefore, we asked if mouse TPCs are sensitive to extracellular ATP. Primary TPC cultures retain high purity for ≥14 days *in vitro* (Figure 1A, and Figure 1–figure supplement 1A&B) and cells express transcripts for several ionotropic (P2X2, P2X4, P2X7) and metabotropic (P2Y2, P2Y6) purinoceptors (Figure 1B). The specific biophysical and pharmacological profile of ATP-dependent transmembrane currents – i.e., saturation at ≤100 μM ATP, modest but persistent desensitization, reduced BzATP sensitivity (data not shown), potentiation by ivermectin, and inhibition by suramin (Figure 1C-G) – strongly suggests functional expression of P2X2 and / or P2X4, but not P2X7 receptors (Coddou et al., 2011). Notably, ~46% of all ATP-sensitive TPCs additionally displayed a delayed, but long-lasting inward current that gradually developed over tens of seconds after ATP stimulation ended (Figure 1–figure supplement 1C&D). Also activated by UTP, dependent on presence of intracellular GTP, and largely carried by Cl^−^ (Figure 1–figure supplement 1E-H), this current likely results from P2Y receptor-mediated phosphoinositide turnover, Ca^2+^ release, and activation of Ca^2+^-gated Cl^−^ channels. Notably, live-cell ratiometric Ca^2+^ imaging in TPCs revealed robust and repetitive cytosolic Ca^2+^ transients upon ATP exposure (Figure 1H&I). Reducing extracellular Ca^2+^ to 100 nM, a concentration approximately equimolar to cytosolic levels, did substantially reduce, but not abolish ATP response amplitudes (Figure 1J&K). The selective P2Y receptor agonist UTP also triggered Ca^2+^ signals (data not shown). Together, these data suggest that mouse TPCs functionally express both ionotropic and metabotropic purinoceptors.

**Figure 1.**
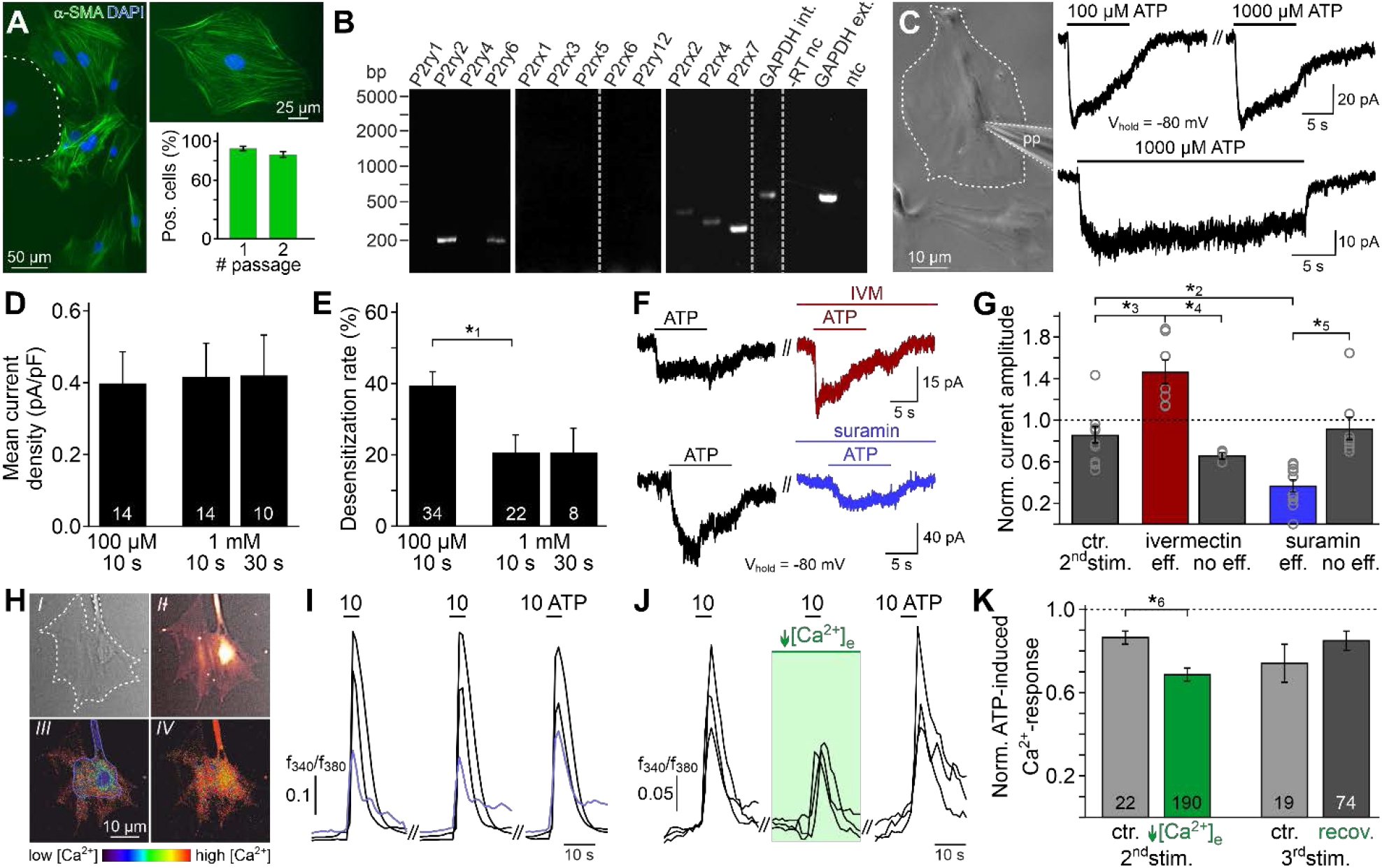
ATP is a potent TPC stimulus. (**A**) Immunostaining against α-smooth muscle actin (α-SMA, green) marks TPCs *in vitro* (Tung and Fritz, 1990). Cell count is determined by nuclear staining (DAPI, blue). Cultures retain high TPC purity for at least two passages (92 ± 2%, n = 1102 (#1); 86 ± 3%, n = 542 (#2)). Dashed line delimits one of the few α-SMA-negative cells. (**B**) RT-PCR profiling of purinoceptor isoforms in TPC cultures reveals P2rx2, P2rx4, P2rx7, P2ry2, and P2ry6 transcripts. Dashed grey vertical lines indicate cuts in a given gel. (**C**-**G**) ATP exposure triggers TPC transmembrane currents. (**C**) Phase-contrast micrograph depicting a TPC (dashed line) targeted by a patch pipette (pp). Original whole-cell recordings illustrate representative currents in response to ATP stimulation (100 μM *versus* 1000 μM and 10 s *versus* 30 s, respectively). (**D**, **E**) Quantification (bar charts; mean ± SEM; n as indicated) reveals saturation of peak current density at ≤100 μM ATP (**D**) and modest desensitization at a concentration-dependent rate (**E**). (**F**) Whole-cell voltage-clamp recordings show ATP-induced currents (100 μM; 10 s) that are potentiated by ivermectin (3 μM) and partially inhibited by suramin (100 μM), respectively (≥60 s preincubation). (**G**) Quantification (bar charts; mean ± SEM; data normalized to initial control response) demonstrates dichotomy in drug sensitivity. (**H**-**K**) ATP-dependent Ca^2+^ mobilization in cultured TPCs. Ca^2+^ transients in response to repetitive stimulation (10 μM, 10 s) are monitored by ratiometric (fura-2) fluorescence imaging. (**H**) Phase contrast (*I*) and merged fluorescence (**f**_380_; *II*) images of a TPC *in vitro*. Bottom pseudocolor frames (rainbow 256 colour map) illustrate relative cytosolic Ca^2+^ concentration ([Ca^2+^]_c_) before (*III*) and during (*IV*) ATP stimulation. (**I**, **J**) Representative original traces from time-lapse fluorescence ratio (*f*_340_/*f*_380_) recordings depict repetitive [Ca^2+^]_c_ elevations upon ATP exposure under control conditions ((**I**) blue traces correspond to the TPC in (**H**)) and during reduced extracellular Ca^2+^ concentration ((**J**) [Ca^2+^]e = 100 nM; 60 s preincubation). (**K**) Bar chart depicting [Ca^2+^]_c_ signal amplitudes (mean ± SEM; n as indicated) – normalized to the initial ATP response – under control conditions (grey) *versus* low [Ca^2+^]_e_ (green). Asterisks denote statistically significant differences (*^1^*p* = 0.001; *^2^*p* = 0.002; *^3^*p* = 5.5e-5; *^4^*p* = 0.0006; *^5^*p* = 0.02; *^6^*p* = 0.02; Student *t*-test (**E**, **K**), one-way ANOVA (**G**)).

Next, we asked if TPCs also exhibit ATP sensitivity in their physiological setting. In parallel approaches, we examined purinergic Ca^2+^ signals from mouse TPCs in acute seminiferous tubule sections (Fleck et al., 2016), using either TPC-specific expression of genetically encoded Ca^2+^ indicators (GCaMP6f) or bulk loading with a synthetic Ca^2+^ sensor (fura-2). First, we confirmed inducible smooth muscle-targeted testicular expression of fluorescent reporter proteins in SMMHC-CreER^T2^ x Ai14D mice (Figure 2A–C, movie S1). Second, TPC-specific GCaMP6f expression in SMMHC-CreER^T2^ x Ai95D mice revealed robust Ca^2+^ transients in cells of the tubular wall upon ATP exposure (Figure 2D&E). Third, fura-2/AM loading of intact seminiferous tubule sections preferentially labelled the outermost cell layer (Figure 2F), allowing semi-quantitative *in situ* imaging of ATP-dependent Ca^2+^ signals in mouse TPCs (Figure 2G&H). So far, our results thus demonstrate that challenging TPCs with extracellular ATP triggers robust Ca^2+^ signals both *in vitro* and *in situ*.

**Figure 2:**
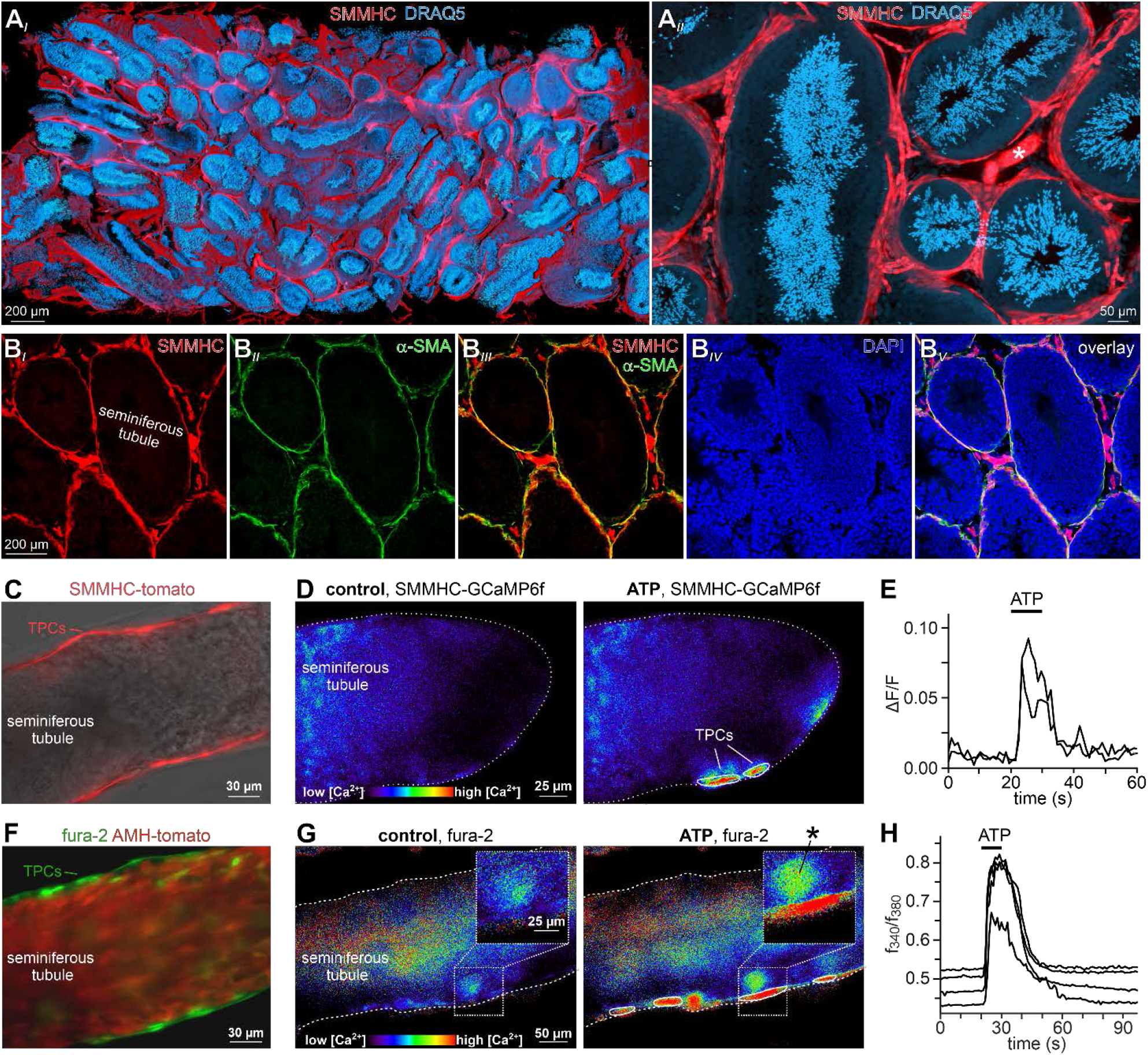
ATP triggers TPC Ca^2+^ signals in acute seminiferous tubule sections. (**A**) 3D reconstruction of an intact 6 × 3 × 1.5 mm testis sample from a SMMHC-CreER^T2^ x Ai14D mouse after tissue clearing (CLARITY (Chung and Deisseroth, 2013)) reveals tdTomato expression (red) restricted to TPCs and endothelial cells (asterisk in (**A**_*II*_)). Nuclear staining (DRAQ5; blue) prominently labels condensed DNA in post-meiotic germ cells. (**B**) SMMHC-CreER^T2^-expressing cells in the tubule wall are TPCs. In testis cryosections from adult SMMHC-CreER^T2^ x Ai14D mice, Cre-driven tdTomato signals (**B**_*I*_) and α-SMA immunostaining (**B**_*II*_) colocalize at tubular margins (**B**_*III*_). Note that endothelial vasculature in interstitial regions is α-SMA-negative (**B**_*V*_). (**C**–**H**) Both TPC-specific expression of a genetically encoded Ca^2+^ indicator (GCaMP6f) and bulk loading with a synthetic Ca^2+^ sensor (fura-2) allow for TPC-selective live cell Ca^2+^ imaging in acute seminiferous tubule sections. (**C**) Merged fluorescence and reflected light micrographs show the location of SMMHC-expressing TPCs (red) within the wall of an intact tubule. (**D**–**E**) Cre-dependent GCaMP6f expression in SMMHC-CreER^T2^ x Ai95D mice reveals Ca^2+^ transients in TPCs in response to ATP. Representative fluorescence images ((**D**) rainbow 256 colour map) before and during ATP exposure (100 μM; 10 s), and corresponding traces (**E**) showing changes in GCaMP6f intensity (ΔF/F) over time. Traces from ROIs outlined in (**D**) (white solid lines). (**F**) Merged fluorescence image of an acute seminiferous tubule section from an AMH-Cre x Ai14D mouse after bulk loading with fura-2/AM (green). Anti-Müllerian hormone (AMH) dependent expression of tdTomato (red) specifically labels Sertoli cells that build the seminiferous epithelium. Note the narrow green band of marginal TPCs that are preferentially labelled by the Ca^2+^-sensitive dye. (**G**–**H**) Ratiometric Ca^2+^ imaging in fura-2-loaded tubules enables semi-quantitative live-cell monitoring of TPC activity. Representative fluorescence images ((**G**) rainbow 256 colour map) before and during ATP exposure (100 μM; 10 s). Corresponding traces (**H**) show the integrated fluorescence intensity ratio (*f*_340_/*f*_380_) from four ROIs (in (**G**); white solid lines) over time. Inset (**G**) shows a putative TPC and an adjacent putative spermatogonium (asterisk) at higher magnification.

### ATP triggers seminiferous tubule contractions

We hypothesized that ATP-induced Ca^2+^ signals in TPCs could mediate contractile motion of the seminiferous tubule. To address this, we established a fast, quasi-simultaneous image acquisition method that enables parallel recording of both peritubular Ca^2+^ responses and seminiferous tubule movement (methods). Brief ATP exposure resulted in a peripheral band of Ca^2+^ activity at the edge of the tubule. Such signals usually coincided with a pronounced contractile motion of the seminiferous tubule (Figure 3A, movie S2). When movement is quantified as the time-lapse image flow field strength (methods) tubular contraction follows the Ca^2+^ signal onset with minimal delay, outlasts the Ca^2+^ signal peak, and recovers slowly (Figure 3B). Both Ca^2+^ responses and tubular movement are dose-dependent and share an ATP threshold concentration of approximately 1 μM (Figure 3C-E, movie S3). Notably, in some tubules, we observed spontaneous low-amplitude ‘vibratory’ movements and local indentations (Figure 4A), reminiscent of the relatively high frequency rippling previously described (Ellis et al., 1981; Worley et al., 1985).

**Figure 3.**
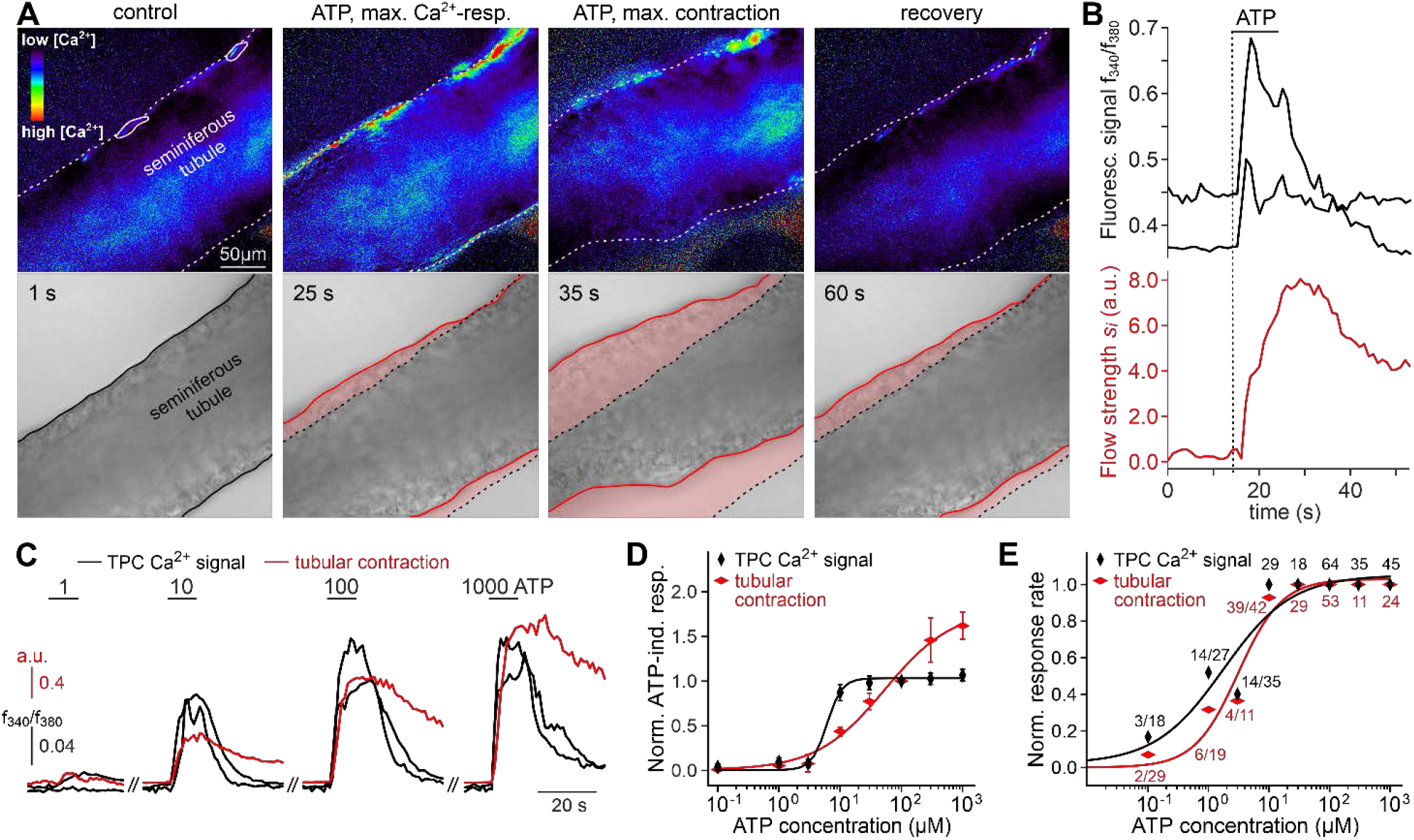
ATP triggers seminiferous tubule contractions. (**A**) Quasi-simultaneous imaging of [Ca^2+^]_c_− dependent fluorescence (top; *f*_340_/*f*_380_; rainbow 256 colour map) and tubular position (bottom; reflected light microscopy). Focus adjusted to provide sharp images of the seminiferous tubule’s edges. Individual frames correspond to the time points indicated, i.e. before, during, and after ATP exposure (see (**B**)). Dashed white lines (top) and corresponding solid black / red lines (bottom) depict the outline of the tubule in each image, respectively. Dotted black lines (bottom) show the contour at t = 1 s for comparison. Pink shades (bottom) accentuate areas that moved. (**B**) Integrated fluorescence ratio (top; black traces correspond to regions-of-interest delimited by solid white lines in (**A**)) and integrated flow strength *s*_*i*_ (bottom; red trace) over time. ATP (100 μM) stimulation as indicated (horizontal bar). With the t = 0 s image as reference, flow strength *s*_*i*_ is calculated by custom code as the average whole tubule pixel shift vector (methods). Dashed vertical line marks the Ca^2+^ signal onset. (**C**–**E**) Ca^2+^ responses and tubular movement are dose-dependent. (**C**) Original traces depict [Ca^2+^]_c_ (black) and tubule movement (red) from a representative experiment. Data calculated as in (**B**). Brief (10 s) stimulations with increasing ATP concentrations (1–1000 μM) trigger dose-dependent Ca^2+^ transients and corresponding contractions. (**D**, **E**) Data quantification. Dose-response curves illustrate peak signals (**D**) and the percentage of responding cells / tubules (**E**). Data are normalized to response to 100 μM ATP (n as indicated in (**E**)).

**Figure 4:**
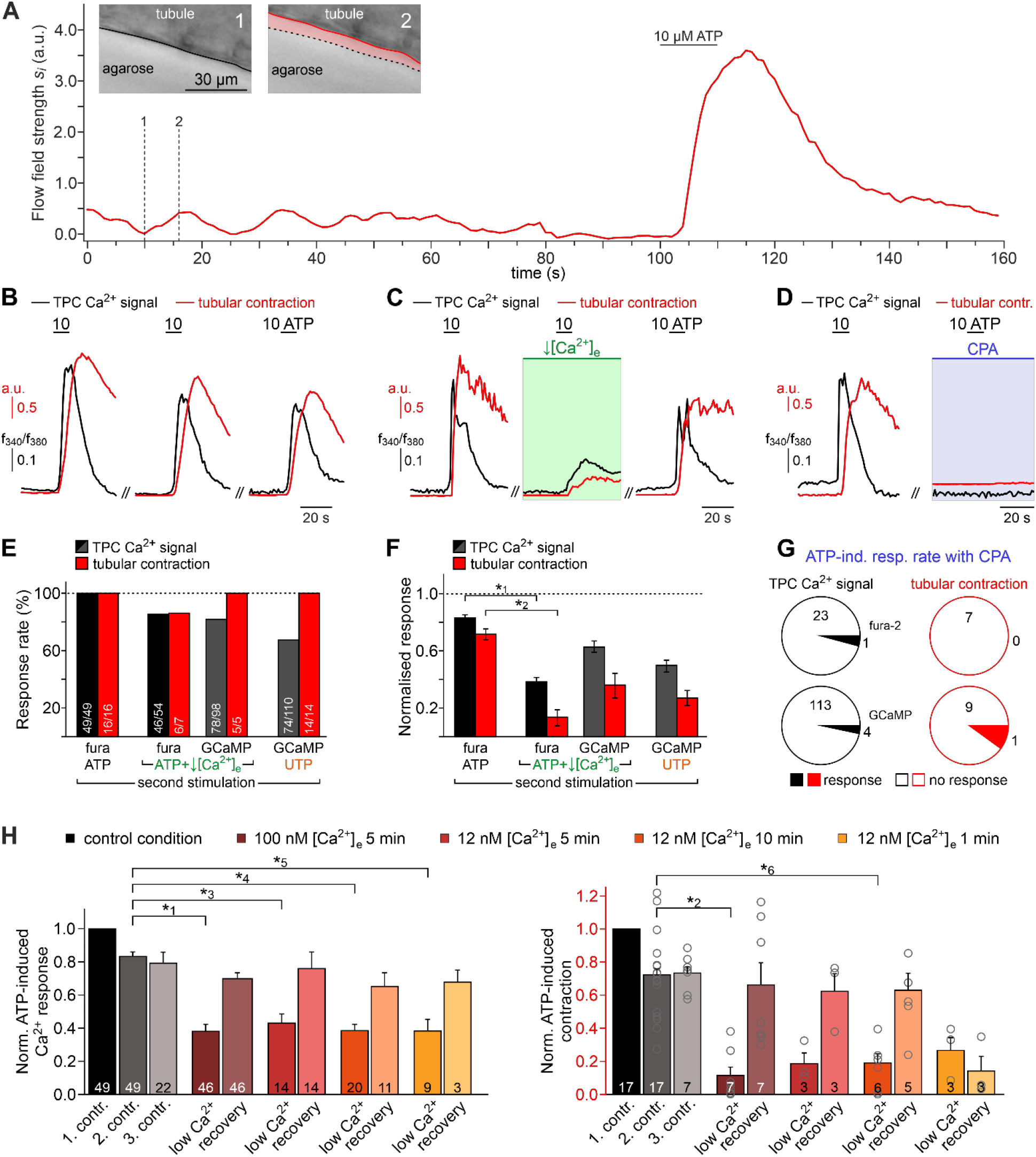
Both intra- and extracellular Ca^2+^ sources contribute to ATP-dependent TPC contractions. (**A**) *In situ* imaging identifies spontaneous low-amplitude ‘vibratory’ movements in acute seminiferous tubule slices. Representative trace illustrating flow field strength analysis of tubular motion under control conditions and upon ATP exposure (10 μM; 10 s). Note that spontaneous indentations share small amplitudes and are restricted to the tubule edge (inset). Black / red lines (inset) depict the outline of the tubule in each image, respectively. Dotted black lines show the contour at t = 1 for comparison. Pink shades accentuate areas that moved. (**B**) Repeated purinergic stimulation triggers robust Ca^2+^ elevations and concurring seminiferous tubule contractions with only minor response adaptation. Traces depict changes in fura-2 intensity ratio (*f*_340_/*f*_380_; black) or tubular movement (red) upon brief ATP exposure (10 s; 10 μM; 5 min intervals) under control conditions. (**C**) Reducing [Ca^2+^]_e_ (100 nM; 5 min incubation) strongly diminishes responses to ATP (10 s; 10 μM). (**D**) Depletion of internal Ca^2+^ stores (CPA (Seidler et al., 1989); 90 μM; ≥15 min incubation) essentially abolishes both Ca^2+^ signals and tubule contractions. (**E**–**G**) Quantification of data exemplified in (**B**–**D**). (**E**) Bar chart depicting ATP sensitivity (response rate; %), independent of signal strength. Occurrence of Ca^2+^ elevations (black) and tubule contractions (red) are plotted for different experimental conditions [i.e., stimulation with ATP or UTP, regular or reduced [Ca^2+^]_e_ (1 mM or 100 nM, respectively), and Ca^2+^ indicator (fura-2 or GCaMP6f, respectively)]. Numbers of experiments are indicated in each bar. (**F**) Signal amplitudes (Ca^2+^, black; contractions, red) of responding TPCs / tubules. Data (mean ± SEM) are normalized to the respective initial response under control conditions (dotted horizontal line). Experimental conditions and numbers of experiments as in (**E**). Asterisks denote statistical significance (*^1^*p* = 2.2e-19 and *^2^*p* = 5.6e^−8^; *t*-test; note: tests only performed when n > 5 and only one variable was changed). (**G**) Pie charts illustrating the profoundly reduced ATP sensitivity of TPCs / tubules after depletion of Ca^2+^ storage organelles (CPA; 90 μM; ≥15 min incubation). (**H**) Effects of lowering [Ca^2+^]_e_ are comparable over both incubation periods and concentrations in the nanomolar range. Significantly reduced, though not abolished TPC Ca^2+^ signals (left) and seminiferous tubule contractions (right) are observed in presence of both 100 nM and 12 nM [Ca^2+^]_e_ as well as for variable incubation periods lasting between 1 and 10 min, respectively. Asterisks denote statistical significance (*^3^*p* = 2.8e^−10^, *^4^*p* = 7.8e^−14^, *^5^*p* = 3.8e^−9^, *^6^*p* = 1.0e^−6^; one-way ANOVA with *post-hoc* Tukey HSD test; note: tests only performed when n ≥ 5).

We next investigated the Ca^2+^ signaling mechanism(s) underlying ATP-dependent TPC contractions. First, we asked whether influx of external Ca^2+^ is involved in TPC force generation. Similar to *in vitro* observations (Figure 1J&K), diminishing or even reversing the driving force for transmembrane Ca^2+^ flux by reducing the extracellular Ca^2+^ concentration to 100 nM or 12 nM, respectively, for variable durations, significantly decreased both TPC Ca^2+^ signals and tubular contractions (Figure 4B-H). These effects were fully reversible. Second, we examined a potential role of ATP-induced Ca^2+^ release from internal storage organelles. Ca^2+^ depletion of the sarcoplasmic reticulum via pharmacological inhibition of the sarco / endoplasmic reticulum Ca^2+^-ATPase essentially abolished both ATP-dependent Ca^2+^ signals and contractions (Figure 4D). Importantly, all results from ratiometric fura-2 imaging were qualitatively indistinguishable from those obtained with genetically targeted GCaMP6f (Figure 4E-G). Third, we aimed to quantify the specific contribution of metabotropic purinoceptors to the overall ATP-mediated effect. The P2Y receptor-selective agonist UTP evoked both TPC Ca^2+^ signals and tubular contractions (Figure 4E&F). However, these responses were substantially reduced compared to control ATP stimulations (Figure 4F), similar to the diminished signals observed under low extracellular Ca^2+^ conditions.

Together, these data strongly suggest that (*i*) extracellular ATP acts as a potent TPC stimulus that triggers seminiferous tubule contractions *in situ*, that (*ii*) both P2X and P2Y receptors mediate TPC responses to ATP exposure, that (*iii*) Ca^2+^ mobilization from the sarcoplasmic reticulum is necessary to evoke TPC responses, and consequently that (*iv*) influx of external Ca^2+^ via ionotropic P2X receptors is not sufficient to drive TPC signals and evoke contractions. Notably, our general finding of ATP-induced mouse TPC contractions is likely transferable to human peritubular cells. When primary human TPC cultures (Walenta et al., 2018) were exposed to extracellular ATP, morphological changes were observed within seconds-to-minutes (Figure 4–figure supplement 1A&B). Moreover, embedding cells in collagen gel lattices revealed considerable contractile force in response to ATP (Figure 4–figure supplement 1C&D).

### ATP drives directional luminal transport

We hypothesized that ATP-induced tubular contractions could impact the transport of luminal fluid and spermatozoa. To test this, we custom-built a whole-mount macroscopic imaging platform, designed to allow both widefield and fluorescence time-lapse imaging of intact seminiferous tubules (Figure 5A). In addition, this setup enables visual categorization of the twelve seminiferous epithelium cycle stages into three distinct stage groups (Hess and De Franca, 2008) and allows precisely timed focal perfusion (methods). First, we asked if brief focal purinergic stimulation triggers seminiferous tubule contractions and, consequently, luminal content movement. Flow field change analysis reveals some basal luminal motion independent of mechanical stimulation (Figure 5B). However, ATP exposure triggered a strong increase in luminal flow field strength that outlasted the presence of ATP for several tens of seconds (Figure 5B, movie S4). Second, we analyzed if luminal movement depends on the tubule’s cycle stage and, consequently, luminal sperm count. When we compared ATP-induced movement between directly stimulated stages (i.e., region-of-interest (ROI) 0) with a high (group II) *versus* a relatively low (groups I & III) amount of luminal sperm, we observed no difference in stimulation-dependent motion (Figure 5C_*I*_). Thus, direct ATP exposure triggers tubular contractions independent of cycle stage and luminal sperm count. Third, we investigated if luminal movement is restricted to the area of stimulation or, by contrast, if motion propagates beyond the directly stimulated tubule section. When we analyzed luminal motion in equidistant tubule sections adjacent to the directly stimulated area (Figure 5A), we found a significant, though relatively small bidirectional wave of propagating movement in stage groups with a low luminal sperm count (Figure 5C_*II*_). Strikingly, we observed strong unidirectional movement upon ATP stimulation of tubule sections with high luminal sperm density (Figure 5C_*II*_). In this stage group, luminal content is predominantly propelled towards areas of ascending spermatogenic cycle stages. These findings demonstrate directionality of sperm transport upon purinergic TPC stimulation.

**Figure 5.**
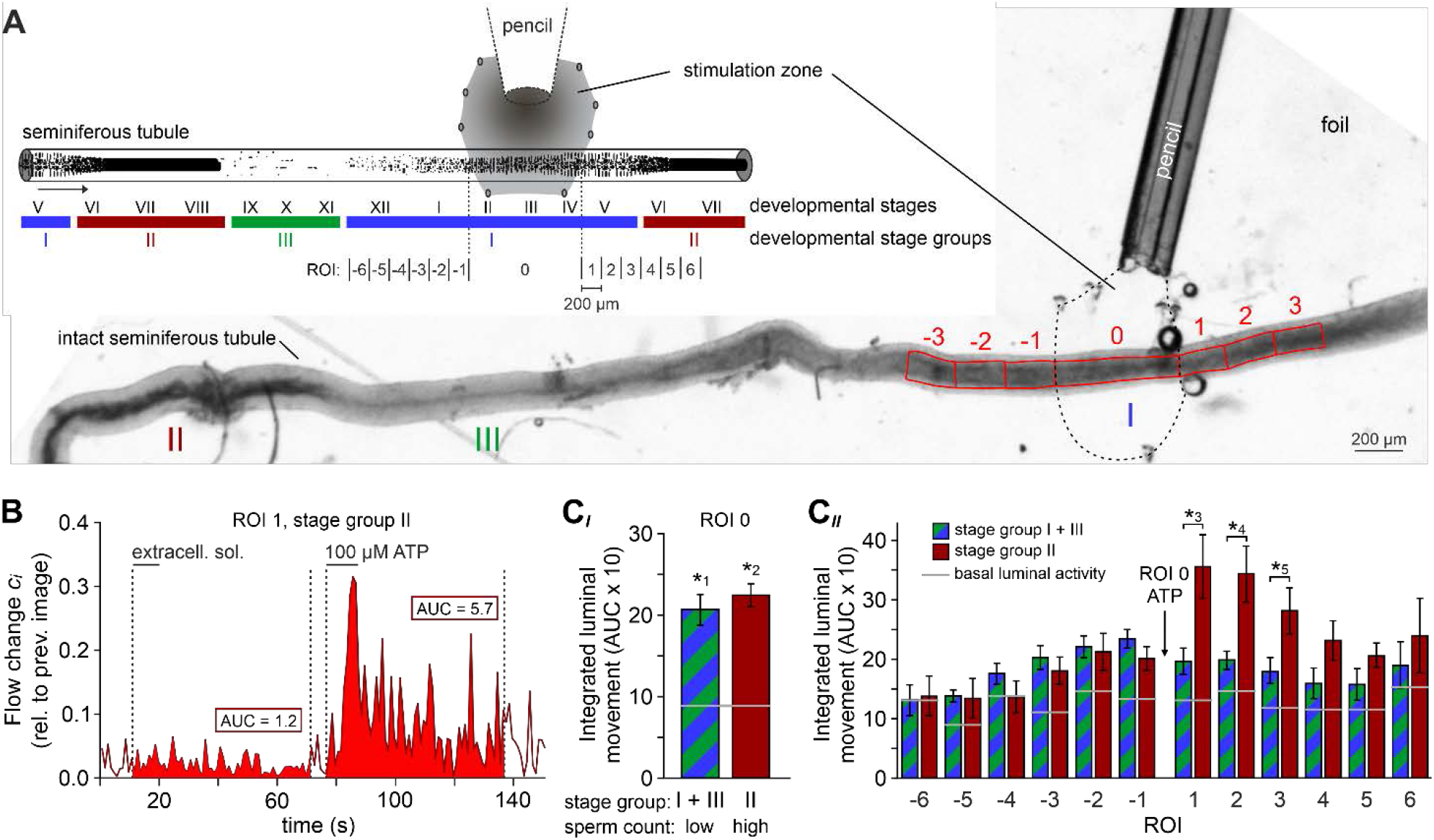
ATP drives directional luminal transport. (**A**) Schematic drawing (top) and original low-magnification image (bottom) of the experimental setup. Intact seminiferous tubules are placed on transparent foil in a custom-built macroscopic imaging chamber. The tubule is kept stationary by gentle suction through tiny holes punched in the foil and vacuum underneath. As previously suggested (Hess and De Franca, 2008), tubules are coarsely categorized into three stages (I–III; color code) according to luminal sperm content. Precise mapping of stimulated regions is feasible by positioning both tubule and perfusion pencil within an area delimited by several holes that outline a stimulation zone (methods). The tubule region directly exposed to ATP is designated as ROI 0, with adjacent equidistant sections numbered consecutively (up to ROI ±6). (**B**) Analysis of luminal content movement by calculation of flow change *c*_*i*_ relative to each previous image (methods) within a representative luminal ROI. Motion is quantified by measuring the area under curve (AUC; solid red) within 60 s after stimulation onset. Note that mechanical control stimulation (extracellular solution) does not affect basal luminal motion. (**C**) Bar charts depicting luminal content movement (means ± SEM) upon ATP stimulation (100 μM; 10 s) in either directly exposed regions (c_*I*_; n = 17) or adjacent areas (c_*II*_; n = 3–17). Green / blue (groups I & III) and red (group II) bars depict stages with a low *versus* a high luminal sperm count, respectively. Horizontal grey lines mark the average basal luminal motion prior to stimulation. ATP induces significantly increased content movement in directly stimulated areas (ROI 0) independent of luminal sperm count / stage group (c_*I*_). Note that in adjacent regions (c_*II*_) unidirectional movement occurs exclusively in tubule sections with high luminal sperm density. Asterisks denote statistically significant differences (*^1^*p* = 8.7e^−5^; *^2^*p* = 6.7e^−7^; *^3^*p* = 0.005; *^4^*p* = 0.002; *^5^*p* = 0.03; unpaired two-tailed *t*-test).

As expected, ATP-induced tubule contractions also manifest as Ca^2+^ signals in TPCs (Figure 6A&B, movie S5). Surprisingly, however, these Ca^2+^ elevations are limited to those areas directly exposed to ATP. We observed no such signals in adjacent tubule sections independent of the stimulated stage group and an ascending or descending stage direction (Figure 6C&D). This finding indicates that ATP acts as a local paracrine messenger that, by itself, is not sufficient to trigger a signal that propagates in a regenerative wave-like fashion along a tubule’s longitudinal axis. In turn, transport directionality likely results from morphological characteristics rather than peristaltic contractility.

**Figure 6:**
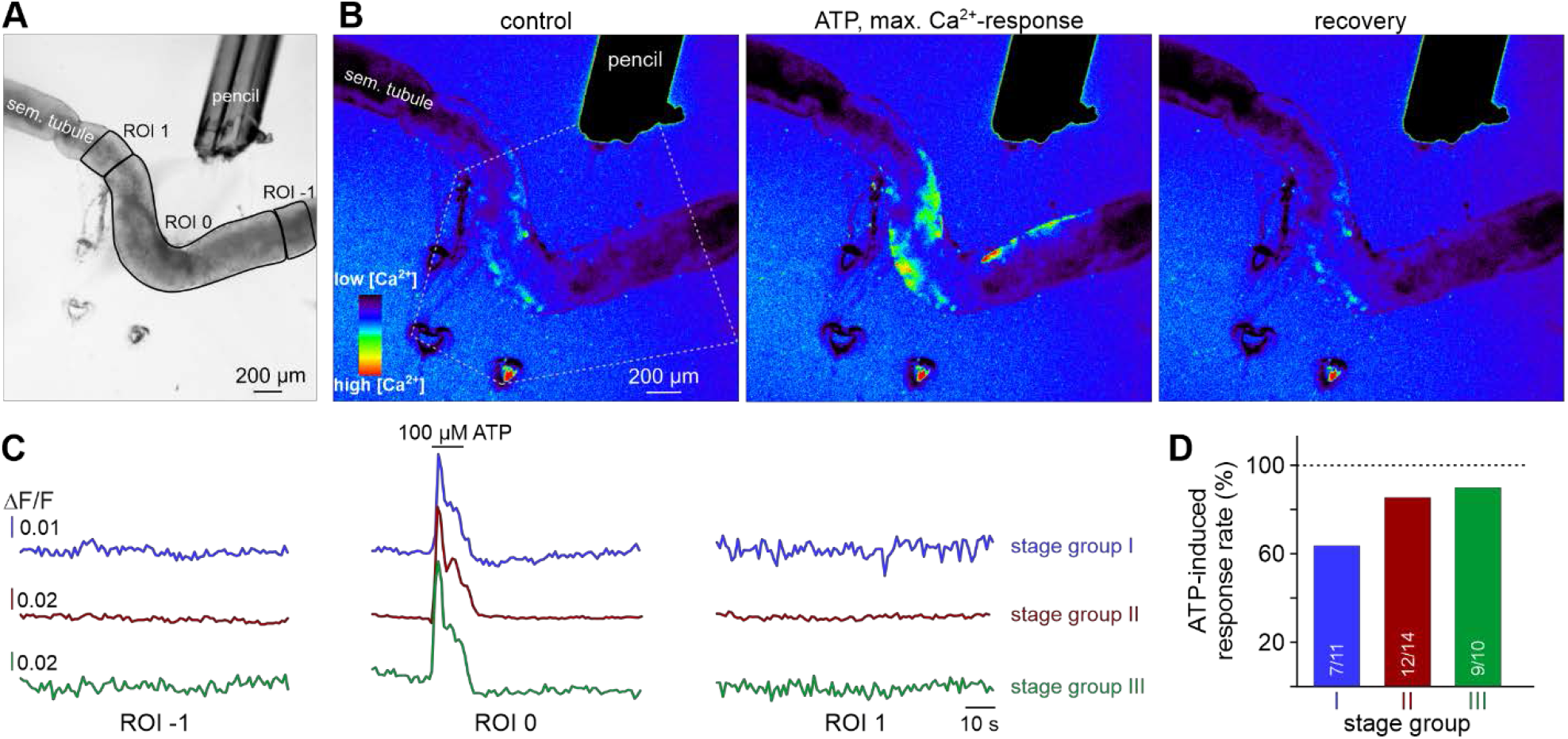
ATP causes Ca^2+^ elevations within a restricted paracrine radius. (**A**) Low-magnification brightfield image of an intact seminiferous tubule segment dissected from SMMHC-CreER^T2^ x Ai95D mice and positioned directly in front of the tip of a 250 μm diameter perfusion pencil. ROIs (black lines) are drawn to encompass the area that is directly exposed to fluid flow (ROI 0) as well as adjacent regions (ROIs 1 and −1), respectively. Suction produced by negative pressure (applied through holes in the elastic foil pad beneath the tubule) limits the area of perfusion. (**B**) Pseudocolor GCaMP6f fluorescence intensity images of the tubule shown in (**A**) reveals Ca^2+^ transients in TPCs in response to ATP. Representative images (rainbow 256 colour map) correspond to time points before, during, and after focal ATP exposure (100 μM; 10 s). The area directly challenged with ATP is denoted by the white dotted lines. For clarity, autofluorescence of the perfusion pencil was removed. Note that Ca^2+^ elevations are limited to ROI 0. (**C**) Representative original recordings of changes in GCaMP6f intensity (ΔF/F) over time from tubule segments of the three different stage groups (I–III). Traces exemplify Ca^2+^ signals (or the lack thereof) in ROIs 0, −1, and 1, respectively. Independent of the epithelial cycle stage investigated, ATP-induced Ca^2+^ elevations are restricted to directly exposed tissue segments. (**D**) Quantification of ATP sensitivity among tubule segments of different cycle stage. Bar charts illustrate that purinergic stimulation causes Ca^2+^ signals irrespective of stage and, thus, luminal sperm count. Numbers of experiments as indicated in bars.

### ATP induces tubular contractions *in vivo*

To ultimately attribute a physiological role to ATP-dependent Ca^2+^ signals in TPCs, tubular contractions, and corresponding transport of luminal content, these phenomena must (*i*) occur spontaneously in living animals, and must (*ii*) be triggered experimentally by ATP exposure *in vivo*. Thus, to investigate any *in vivo* relevance of our findings, we designed a custom-built 3D printed *in vivo* imaging stage (Figure 7–figure supplement 1) that allows both widefield epi-fluorescence and multiphoton microscopy of the mouse testis.

Initially, we monitored spontaneous seminiferous tubule activity in SMMHC-CreER^T2^ x Ai95D mice. Multiphoton time-lapse imaging revealed spontaneous TPC Ca^2+^ signals that typically accompanied strong tubule contractions (Figure 7A&B, movie S6). Several characteristics emerged from quantitative analysis of these observations. First, during sufficiently long recording periods (≤10 min), contractions occur in essentially all seminiferous tubules (Figure 7–figure supplement 2A). Second, contractions of individual tubules are not synchronized (Figure 7B). Third, periods of enhanced activity (≥2 contractions within 90 s) are interrupted by long episodes of quiescence (Figure 7B, Figure 7–figure supplement 2B). Fourth, the durations of TPC Ca^2+^ signals and corresponding contractions are positively correlated (Figure 7–figure supplement 2C), confirming a causal relationship.

**Figure 7.**
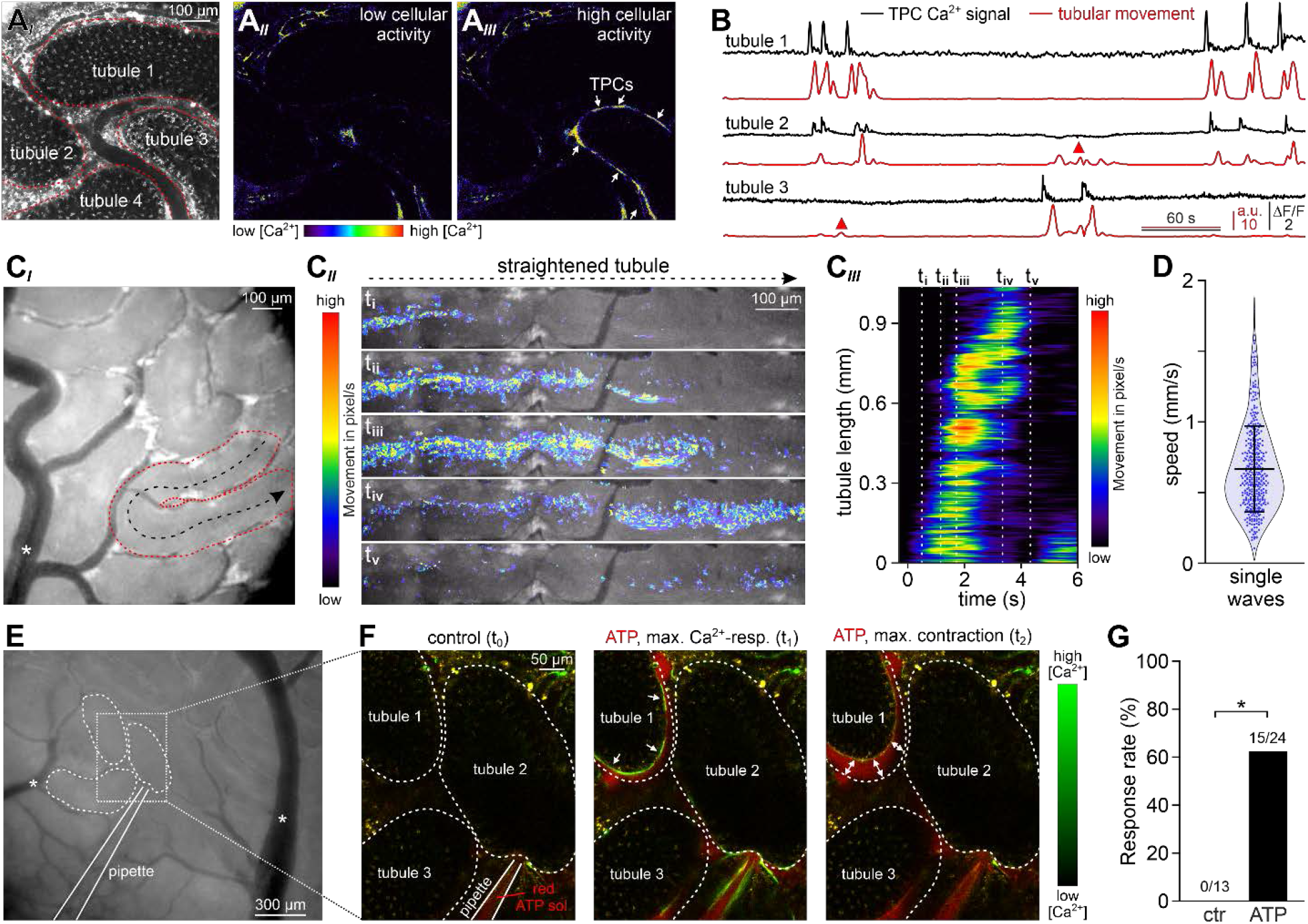
ATP induces tubular contractions *in vivo*. (**A**) Multiphoton *in vivo* fluorescence microscopy in SMMHC-CreER^T2^ x Ai95D mice enables time-lapse imaging of TPC activity. Maximum grey scale projection outlines segments from four seminiferous tubules (red dotted lines (**A**_*I*_)). Pseudocolor images of GCaMP6f intensity indicate [Ca^2+^]_c_ changes in TPCs of tubule 3 during phases of low (**A**_*II*_) *versus* high (**A**_*III*_) spontaneous activity (rainbow colour map; white arrows in (**A**_*III*_)). (**B**) Original traces depict simultaneous TPC Ca^2+^ signals (black; ΔF/F) and tubular contractions (red; calculated as flow change *c*_*i*_ relative to each previous image (methods) over time in tubules 1–3 (**A**). Red triangles mark passive movements, which occur upon contractions of adjacent tubules. (**C**) Analysis of spontaneous tubular motion *in vivo*. Low magnification incident light image of the mouse testis (**C**_*I*_) shows several superficial seminiferous tubule segments, testicular blood vessels (white asterisk; note that unobstructed blood supply (i.e., visualizing erythrocyte flow) is checked routinely), and a specific segment outlined by red dotted lines. After time-lapse imaging, this segment is digitally straightened (**C**_*II*_) and subjected to motion analysis. For different time points (**i**–**v**), pixel movement and its propagation are reflected by merged pseudocolor images. Directionality is indicated by the black arrow in (**C**_*I*_). From a kymograph (**C**_*III*_), the time–space relationship of tubular motion becomes apparent (time points **i**–**v** as indicated by dashed vertical lines). (**D**) Violin plot depicting the velocity of contractile movement in individual tubule segments (blue dots). (**E**–**G**) ATP-induced Ca^2+^ signals and contractions *in vivo*. (**E**) Low magnification epi-fluorescence image of several superficial seminiferous tubule segments and blood vessels (white asterisks). The boxed area includes three tubule segments (dotted black lines), which are targeted by a low resistance pipette filled with fluorescently labelled ATP solution. (**F**) Enlarged view of the area outlined in (**E**). Merged (red / green) multiphoton fluorescence images taken before and during / after brief stimulation with ATP. The middle and right frames correspond to the point of maximum Ca^2+^ signal (green) and contraction (double arrows) of tubule 1, respectively. (**G**) Bar chart quantification of contractions induced by nanoliter puffs of saline with or without ATP (1 mM). Asterisk denotes statistical significance (*p* = 0.036; Fisher’s Exact test).

Next, we asked whether spontaneous *in vivo* contractions are coordinated along the longitudinal tubular axis. Low magnification incident light microscopy enabled simultaneous observation of several superficial seminiferous tubule segments (Figure 7C). Movement analysis along the length of digitally straightened tubules demonstrates wave-like unidirectional motions that propagate with high velocities (Figure 7C&D). Notably, these coordinated contractile movements provide sufficient force to ensure luminal sperm transport (movie S7).

Finally, we examined if brief focal ATP stimulation also triggers peritubular Ca^2+^ signals and seminiferous tubule contractions *in vivo*. Therefore, we filled low resistance patch pipettes with fluorescently labelled ATP solution, penetrated the *tunica albuginea*, and targeted the interstitial space close to neighboring tubules (Figure 7E, movie S8). Nanoliter puffs of ATP-containing test solution induced both Ca^2+^ transients in genetically labeled TPCs and strong tubule contractions in the majority of experiments (Figure 7F&G). By contrast, puffs of extracellular saline rarely stimulated any such response (Figure 7–figure supplement 2D). Taken together, *in vivo* recordings demonstrate that robust recurrent seminiferous tubule contractions (*i*) occur spontaneously, (*ii*) are driven by cytosolic Ca^2+^ elevations in TPCs, and (*iii*) can be triggered experimentally by ATP exposure. Consequently, paracrine purinergic signaling in the mouse testis is a mediator of luminal sperm transport within the seminiferous tubule network.

## Discussion

The molecular and cellular mechanisms that control paracrine testicular communication have to a large extent remained controversial, if not elusive (Schlatt and Ehmcke, 2014). For TPCs in particular, a contractile function under paracrine control and, consequently, a critical role in male infertility have long been proposed (Albrecht et al., 2006; Romano et al., 2005), but direct experimental evidence has been lacking (Mayerhofer, 2013). While several signaling molecules, including vasopressin (Pickering et al., 1989), oxytocin (Worley et al., 1985), prostaglandins (Hargrove et al., 1975), endothelin (Filippini et al., 1993), and others (Albrecht et al., 2006; Mayerhofer, 2013), have been proposed to act on TPCs, a role of ATP in seminiferous tubule contractility has been explicitly ruled out early on (Hovatta, 1972). By contrast, our data reveal ATP is a strong stimulus that activates TPCs via P2X and P2Y receptors, mediating coordinated tubule contractions and luminal sperm transport *in situ* and *in vivo*. Both spontaneous and ATP-dependent contractions trigger fast, stage-dependent, and directional transport of luminal content. It is thus tempting to speculate that seminiferous tubule contractility in general, and purinergic TPC signaling in particular, are promising targets for male infertility treatment and / or contraceptive development.

## Materials and Methods

### Animals

All animal procedures were approved by local authorities and in compliance with both European Union legislation (Directive 2010/63/EU) and recommendations by the Federation of European Laboratory Animal Science Associations (FELASA). When possible, mice were housed in littermate groups of both sexes (room temperature (RT); 12:12 h light-dark cycle; food and water available *ad libitum*). If not stated otherwise, experiments used adult (>12 weeks) males. Mice were killed by CO_2_ asphyxiation and decapitation using sharp surgical scissors. We used C57BL/6J mice (Charles River Laboratories, Sulzfeld, Germany) as well as offspring from crossing either SMMHC-CreER^T2^ (JAX #019079) (Wirth et al., 2008) or 129S.FVB-Tg(Amh-cre)8815Reb/J (JAX #007915) (Holdcraft and Braun, 2004) mice with either Ai95D (JAX #028865) (Madisen et al., 2015) or Ai14D (JAX #007914) (Madisen et al., 2010) mice, respectively.

### Chemicals and solutions

The following solutions were used:

(**S**_**1**_) 4-(2-Hydroxyethyl)piperazine-1-ethanesulfonic acid (HEPES) buffered extracellular solution containing (in mM) 145 NaCl, 5 KCl, 1 CaCl_2_, 0.5 MgCl_2_, 10 HEPES; pH = 7.3 (adjusted with NaOH); osmolarity = 300 mOsm (adjusted with glucose).
(**S**_**2**_) Oxygenated (95% O_2_, 5% CO_2_) extracellular solution containing (in mM) 120 NaCl, 25 NaHCO^3^, 5 KCl, 1 CaCl_2_, 0.5 MgCl_2_, 5 N,N-bis(2-hydroxyethyl)-2-aminoethanesulfonic acid (BES); pH = 7.3; 300 mOsm (glucose).
(**S**_**3**_) Extracellular low Ca^2+^ solution containing (in mM) 145 NaCl, 5 KCl, 0.5 MgCl_2_, 10 HEPES; pH = 7.3 (NaOH); osmolarity = 300 mOsm (glucose); [Ca^2+^]_free_ = ~110 nM (1 mM EGTA, 0.5 mM CaCl_2_) or 12 nM (1 mM EGTA, 0.1 mM CaCl_2_).
(**S**_**4**_) Oxygenated (95% O_2_, 5% CO_2_) extracellular solution containing (in mM) 120 NaCl, 25 NaHCO^3^, 5 KCl, 0.5 MgCl_2_, 5 BES; pH = 7.3; 300 mOsm (glucose); [Ca^2+^]_free_ = 110 nM (1 mM EGTA, 0.5 mM CaCl_2_) or 12 nM (1 mM EGTA, 0.1 mM CaCl_2_).
(**S**_**5**_) Gluconate-based extracellular solution containing (in mM) 122.4 Na gluconate, 22.6 NaCl, 5 KCl, 1 CaCl_2_, 0.5 MgCl_2_, 10 HEPES; pH = 7.3 (adjusted with NaOH); osmolarity = 300 mOsm (glucose).
(**S**_**6**_) Standard pipette solution containing (in mM) 143 KCl, 2 KOH, 1 EGTA, 0.3 CaCl_2_, 10 HEPES ([Ca^2+^]_free_ = 110 nM); pH = 7.1 (adjusted with KOH); osmolarity = 290 mOsm (glucose).
(**S**_**7**_) Gluconate-based pipette solution containing (in mM) 110 Cs gluconate, 30 CsCl, 2 CsOH, 1 EGTA, 0.3 CaCl_2_, 10 HEPES ([Ca^2+^]_free_ = 110 nM); pH = 7.1 (adjusted with CsOH); osmolarity = 290 mOsm (glucose).

In some experiments Na-GTP (0.5 mM) was added to the pipette solution. Free Ca^2+^ concentrations were calculated using WEBMAXCLITE v1.15. If not stated otherwise, chemicals were purchased from Sigma (Schnelldorf, Germany). Cyclopiazonic-acid (CPA) and 2’(3’)-O-(4-Benzoylbenzoyl)adenosine-5’-triphosphate (BzATP) triethylammonium salt was purchased from Tocris Bioscience (Bristol, UK). Fura-2/AM was purchased from Thermo Fisher Scientific (Waltham, MA). Final solvent concentrations were ≤0.1%. When high ATP concentrations (≥1 mM) were used, pH was readjusted.

### Stimulation

For focal stimulation, solutions and agents were applied from air pressure-driven reservoirs via an 8-in-1 multi-barrel ‘perfusion pencil’ (AutoMate Scientific; Berkeley, CA). Changes in focal superfusion (Veitinger et al., 2011a) were software-controlled and, if required, synchronized with data acquisition by TTL input to 12V DC solenoid valves using a TIB 14S digital output trigger interface (HEKA Elektronik, Lambrecht/Pfalz, Germany). For focal stimulation during *in vivo* recordings, ATP was puffed from pulled glass pipettes using a microinjection dispense system (Picospritizer III; Parker Hannifin, Hollis, NH).

Extracellular low Ca^2+^ solutions (**S**_**3**_ & **S**_**4**_) were applied via both the bath and perfusion pencil. To ensure depletion of Ca^2+^ stores by CPA we monitored intracellular Ca^2+^ levels during drug treatment (0.05 Hz frame rate). Transient CPA-dependent Ca^2+^ elevations lasted 10 – 40 min. After baseline Ca^2+^ levels were restored, cells / slices were again challenged with ATP. Control recordings, omitting CPA, were performed under the same conditions.

### Slice preparation

Acute seminiferous tubule slices were prepared as previously described (Fleck et al., 2016) with minor modifications. Briefly, seminiferous tubules from young adults were isolated after *tunica albuginea* removal, embedded in 4% low-gelling temperature agarose (VWR, Erlangen, Germany), and 250 μm slices were cut with a VT1000S vibratome (Leica Biosystems, Nussloch, Germany). Acute slices were stored in a submerged, oxygenated storage container (**S**_**2**_; RT). When using testicular tissue from Ai95D mice, slices were protected from light during storage to avoid GCamP6f bleaching.

### TPC culture

After mouse testis isolation and removal of the *tunica albuginea,* the seminiferous tubules were placed in Dulbecco’s Modified Eagle Medium / Nutrient Mixture F-12 (DMEM/F-12; Invitrogen) containing 1 mg ml^−1^ collagenase A and 6 μg ml^−1^ DNase (10 min; 34°C; shaking water bath (60 cycles min^−1^)). Three times, the samples were washed (DMEM/F-12; 5 ml), allowed to settle for 5 min, and the supernatant was discarded. Next, tubules were incubated DMEM/F-12 containing 1 mg ml^−1^ trypsin and 20 μg ml^−1^ DNase (20 min; 34°C; shaking water bath (60 cycles min^−1^)). Digestion was stopped by addition of 100 μg ml^−1^ soybean trypsin inhibitor (SBTI) and 20 μg ml^−1^ DNase in phosphate-buffered saline (D-PBS). Then, samples were allowed to settle for 5 min and the supernatant was collected. After two more cycles of washing (DMEM/F-12), settling (5 min), and supernatant collection, the collected cell suspension was centrifuged (10 min; 400*g*) and the supernatant discarded. The pellet was resuspended in DMEM containing FBS (10%) and penicillin G / streptomycin (1%), filtered (cell strainer (100 μm)), and cells were plated in 75 cm^2^ cell culture flask (T75; Invitrogen) and placed in a humidified incubator (37°C; 5% CO_2_). Approximately ⅓ of medium volume was replaced every 3 days. Cells usually reached 100% confluence after 7 days *in vitro* (DIV). Then, cells were washed twice (DPBS^−/−^; 5 min; 37°C) and incubated in 0.05% trypsin / EDTA (5 min; 37°C). Detachment of cells was checked visually and, if necessary, facilitated mechanically. The cell suspension was centrifuged (3 min; 800*g*) and the supernatant discarded. The pellet was resuspended in DMEM at cell densities of ~10^5^ cells ml^−1^ and plated again either in culture flasks or on glass coverslips in 35 mm dishes for experimental use. Again, ⅓ of medium volume was replaced every 3 days. Experiments were performed for ≤5 days after passage.

Human TPCs were isolated from small testicular tissue fragments derived from consenting donors with obstructive azoospermia and normal spermatogenesis as described (Albrecht et al., 2006; Walenta et al., 2018). The study was approved by the local ethical committee (Ethikkommission, School of Medicine, TU Munich, project 169/18S).

### Gene expression analysis

Total RNA was isolated and purified from cultured mouse TPCs (passage 1) with Trizol followed by complementary DNA synthesis with RevertAid™ H Minus kit (#K1632 Thermo Fisher) according to the manufacturer’s instructions. Controls in which the reverse transcriptase was omitted were routinely performed. PCR amplification was performed during 30 thermal cycles (95°C, 20 s; 58°C, 20 s; 72°C, 20 s). The following specific primer pairs were used for PCR amplification:

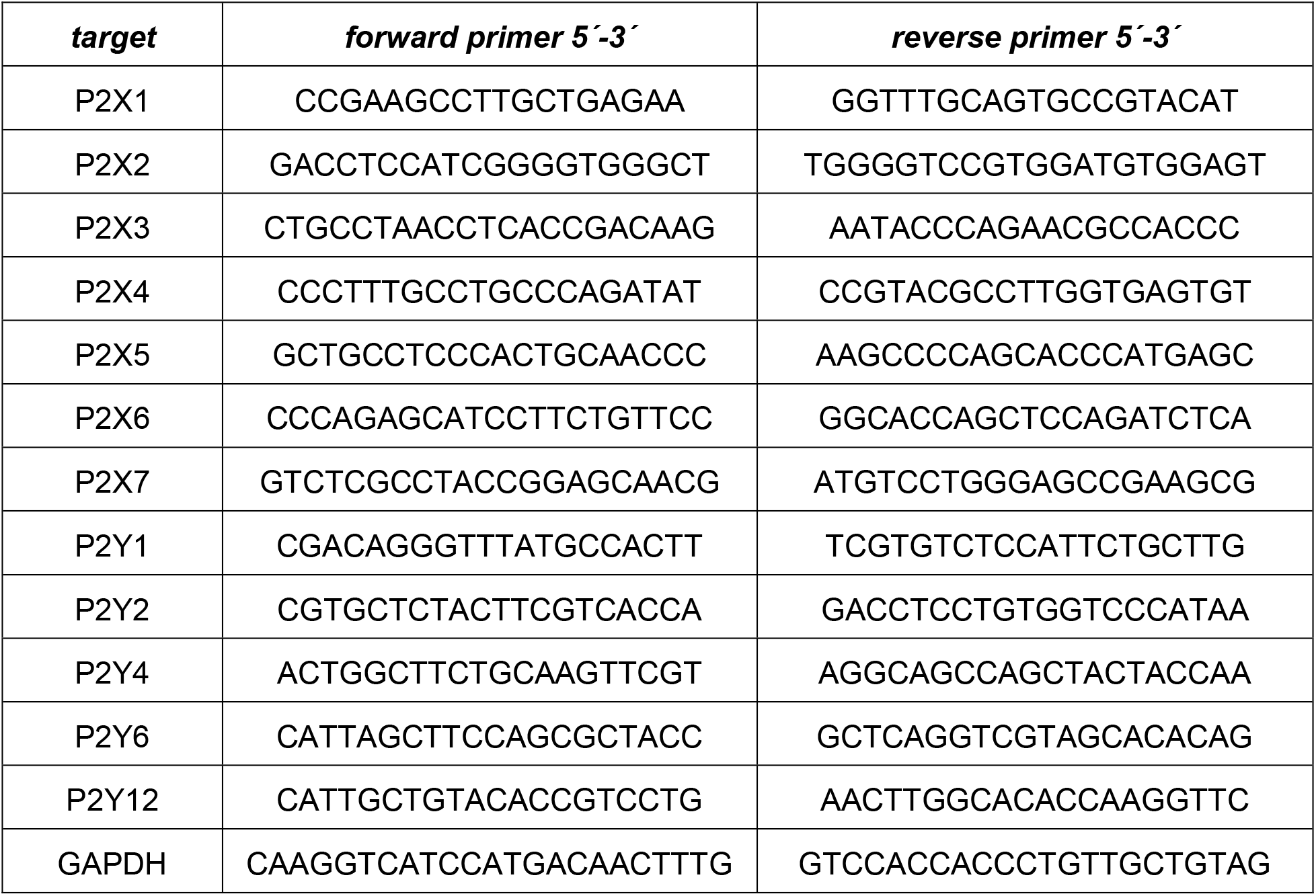

### Immunochemistry and tissue clearing

For immunochemistry of testicular cryosections, testes were fixed with 4% (w/v) paraformaldehyde (PFA) in PBS^−/−^(10 mM, pH 7.4; ≥12 h; 4°C) and subsequently cryoprotected in PBS^−/−^containing 30% sucrose (≥24 h; 4°C). Samples were then embedded in Tissue Freezing Medium (Leica Biosystems), sectioned at 20 μm on a Leica CM1950 cryostat (Leica Biosystems), and mounted on Superfrost Plus slides (Menzel, Braunschweig, Germany). For immunostaining of cultured mouse TPCs, cells were washed (3x; PBS^−/−^), fixed with ice-cold 4% PFA in PBS^−/−^(20 min; RT), and washed again (3x; PBS^−/−^). For blocking, sections / cells were incubated in PBS^−/−^ containing Tween-20 (0.1%) / BSA (3%) solution (1 h; RT). After washing (PBS^−/−^; 2 × 5 min), sections / cells were incubated FITC-conjugated monoclonal anti-actin, α-smooth muscle (α-SMA-FITC, cat # F3777, MilliporeSigma) antibody (1:500 in 3% BSA; 1 h; RT). Excess antibodies were removed by washing (2 × 5 min PBS^−/−^). For nuclear counterstaining, sections / cells were then incubated in PBS^−/−^ containing either DAPI (5 μg ml^−1^; 10 min; RT; Thermo Fisher Scientific) or DRAQ5 (1:500; 5 min; RT; Thermo Fisher Scientific).

Fluorescent images were taken using either an inverted microscope (Leica DMI4000B, Leica Microsystems) or an upright fixed stage scanning confocal microscope (TCS SP5 DM6000 CFS; Leica Microsystems) equipped with a 20x 1.0 NA water immersion objective (HCX APO L; Leica Microsystems). To control for non-specific staining, experiments in which the primary antibody was omitted were performed in parallel with each procedure. Digital images were uniformly adjusted for brightness and contrast using Adobe Photoshop CS6 (Adobe Systems, San Jose, CA, USA).

For testicular tissue clearing we adopted the CLARITY method (Chung et al., 2013) with minor modifications (Gretenkord et al., 2019). Briefly, testes from adult mice were fixed overnight at 4°C in hydrogel fixation solution containing 4% acrylamide, 0.05% bis-acrylamide, 0.25% VA-044 Initiator, 4% PFA in PBS^−/−^ to maintain structural integrity. After hydrogel polymerization, lipids were removed by incubation in 4% sodium dodecyl phosphate (SDS) solution with 200 mM boric acid (pH 8.5) over periods of two months. Solutions were changed bi-weekly. During the final incubation period, the nuclear marker DRAQ5 (1:1000) was added. After washing (2 d) with PBST (0.1% TritonX), samples were incubated for 24 h in RIMS80 containing 80 g Nycodenz, 20 mM PS, 0.1% Tween 20, and 0.01% sodium acid. Cleared samples were imaged using a Leica TCS SP8 DLS confocal microscope, equipped with a digital light-sheet module, 552 nm and 633 nm diode lasers, a HC PL FLUOTAR 5x/0.15 IMM DLS objective (observation), a L 1.6x/0.05 DLS objective (illumination), a DLS TwinFlect 7.8 mm Gly mirror cap, and a DFC9000 sCMOS camera. Rendering and three-dimensional reconstruction of fluorescence images was performed using Imaris 8 microscopy image analysis software (Bitplane, Zurich, Switzerland).

### Electrophysiology

Whole-cell patch-clamp recordings were performed as described (Fleck et al., 2016; Veitinger et al., 2011b). Briefly, mouse TPCs were transferred to the stage of an inverse microscope (DMI 4000B, Leica Microsystems), equipped with phase contrast objectives and a cooled CCD-camera (DFC365FX, Leica Microsystems). Cells were continuously superfused with solution **S**_**1**_ (~3 ml min^−1^; gravity flow; ~23°C). Patch pipettes (~5 MΩ) were pulled from borosilicate glass capillaries with filament (1.50 mm OD / 0.86 mm ID; Science Products) on a PC-10 vertical two-step micropipette puller (Narishige Instruments, Tokyo, Japan), fire-polished (MF-830 Microforge; Narishige Instruments) and filled with **S**_**6**_. An agar bridge (150 mM KCl) connected reference electrode and bath solution. An EPC-10 amplifier controlled by Patchmaster 2.9 software (HEKA Elektronik) was used for data acquisition. We monitored and compensated pipette and membrane capacitance (*C*_mem_) as well as series resistance (R_series_). *C*_mem_ values served as a proxy for the cell surface area and, thus, for normalization of current amplitudes (i.e., current density). Cells displaying unstable R_series_ values were not considered for further analysis. Liquid junction potentials were calculated using JPCalcW software (Barry, 1994) and corrected online. Signals were low-pass filtered [analog 3- and 4-pole Bessel filters (−3 dB); adjusted to ⅓ - ⅕ of the sampling rate (10 kHz)]. Holding potential (*V*_hold_) was −60 mV.

### Fluorescence Ca^2+^ imaging

Cultured mouse TPCs were imaged as described (Veitinger et al., 2011b). Briefly, cells were loaded with fura-2/AM in the dark (5 μM; 30 min; RT; **S**_**1**_) and imaged with an upright microscope (Leica DMI6000FS, Leica Microsystems) equipped for ratiometric live-cell imaging with a 150W xenon arc lamp, a motorized fast-change filter wheel illumination system for multi-wavelength excitation, a CCD camera (DFC365 FX, Leica), and Leica LAS X imaging software. Ten to thirty cells in randomly selected fields of view were viewed at 20x magnification and illuminated sequentially at 340 nm and 380 nm (cycle time 2 s). The average pixel intensity at 510 nm emission within user-selected ROIs was digitized and calculated as the *f*_340_/*f*_380_ intensity ratio.

For parallel recordings of intracellular Ca^2+^ signals and tubular contractions, acute seminiferous tubule slices were bulk-loaded with fura-2/AM in the dark (30 μM; 30 min; RT). After washing (3x; **S**_**1**_), slices were transferred to a recording chamber and imaged with an upright microscope (Leica DMI6000FS, see above). We installed a custom-built reflective shield beneath the recording chamber for parallel monitoring of fluorescence and reflected light. At 1 Hz imaging cycles, we thus recorded two 510 nm fluorescence images (340 / 380 nm excitation) and a ‘pseudo-brightfield’ reflected light image that allowed quasi simultaneous analysis of intracellular Ca^2+^ and tubular movement.

### Whole-mount seminiferous tubule imaging

Isolated tubules (>1 cm length) were placed onto a membrane within a custom-built 3D printed two-compartment recording chamber that was constantly superfused with **S**_**1**_. Small membrane holes under the tubules and around a defined stimulation area allowed for (***i***) gentle fixation of the tubules and (***ii***) focal ATP perfusion of selected tubular regions by vacuum-generated negative pressure (80-180 mmHg) in the submembraneous chamber compartment and continuous suction of **S**_**1**_ from the top compartment. After visual determination of tubular stage (***I*** – ***III***) (Parvinen, 1982), the perfusion pencil was positioned to selectively stimulate an area of known and homogeneous stage. Focal stimulation in the desired area was routinely confirmed by transient dye perfusion (Fast Green) prior to ATP exposure. ATP stimulations (100 μM; 10 s) and corresponding negative controls were compared to determine ATP-dependent Ca^2+^ signals (offspring from crossing SMMHC-CreER^T2^ and Ai95D mice) or tubular contractions and sperm transport. For low-magnification brightfield or fluorescence imaging, we used a MacroFluo Z16 APO A system (Leica Microsystems) equipped with either a DFC450C camera and a PLANAPO 1.0x / WD 97 mm objective (brightfield) or with a monochrome DFC365FX camera and a 5.0x/0.50 LWD PLANAPO objective (fluorescence). Images were acquired at 1 Hz.

### *In vivo* imaging

We administered tamoxifen (75 mg tamoxifen kg^−1^ body weight) to double-positive adult male *offspring* (SMMHC-CreER^T2^ x Ai95D) via daily intraperitoneal injections for 5 consecutive days. Mice were closely monitored for any adverse reactions to the treatment. Experiments were performed 2–5 weeks after the first injection. For surgery, mice were anesthetized with ketamine-xylazine-buprenorphine (100 mg kg^−1^, 10 mg kg^−1^, 0.05–0.1 mg kg^−1^, respectively; Reckitt Benckiser Healthcare, UK). First, we made an incision next to the *linea alba* in the hypogastric region, followed by a 5 mm incision into the peritoneum. One testis was gently lifted from the abdominal cavity. Its *gubernaculum* was cut and the testis – with the spermatic cord, its blood vessels and *vas deferens* still intact – was transferred to a temperature-controlled imaging chamber filled with extracellular solution (**S**_**1**_; 35°C), mounted on custom-designed 3D printed *in vivo* stage (Figure 7–figure supplement 1). Throughout each experiment, vital signs (heartbeat, blood oxygen level, breathing rhythm) were constantly monitored and recorded (breathing). Moreover, we routinely checked unobstructed blood flow within testicular vessels during experiments. To avoid movement artifacts, the tunica was glued to two holding strings using Histoacryl tissue adhesive. After surgery, anesthesia was maintained by constant isoflurane inhalation (1–1.5% in air). Time-lapse intravital imaging was performed using a Leica TCS SP8 MP microscope. For incident light illumination / reflected light widefield recordings (5–10 Hz), we used N PLAN 5x/0.12 or HC APO L10x/0.30 W DLS objectives with large fields of view. Multiphoton time-lapse images were acquired at ~2 Hz frame rates using external hybrid detectors and the HCX IRAPO L25x/0.95 W objective at 930 nm excitation wavelength. For *in vivo* stimulation experiments, we used a Picospritzer III (Parker Hannifin, Pine Brook, NJ) to puff nanoliter volumes of control saline (**S**_**1**_; containing Alexa Fluor 555 (4 μM)) or stimulus solution (**S**_**1**_; containing Alexa Fluor 555 (4 μM) and ATP (1 mM)), respectively, from beveled glass micropipettes onto the surface of seminiferous tubules.

### Data analysis

All data were obtained from independent experiments performed on at least three days. Individual numbers of cells / tubules / experiments (n) are denoted in the respective figures and / or legends. If not stated otherwise, results are presented as means ± SEM. Statistical analyses were performed using paired or unpaired *t*-tests, one-way ANOVA with Tukey’s HSD *post hoc* test or the Fisher Exact test (as dictated by data distribution and experimental design). Tests and corresponding *p*-values that report statistical significance (≤0.05) are individually specified in the legends. Data were analyzed offline using FitMaster 2.9 (HEKA Elektronik), IGOR Pro 8 (WaveMetrics), Excel 2016 (Microsoft, Seattle, WA), and Leica LAS X (Leica Microsystems) software. Dose-response curves were fitted by the Hill-equation. Time-lapse live-cell imaging data displaying both Ca^2+^ signals and tubular contractions were analyzed using custom-written code in MATLAB (The MathWorks, Natick, MA).

For quantitative image analysis, images from both reflected light and fluorescence time-lapse recordings were registered to their respective first image frame at time point t_0_, using the registration algorithm from (Liu et al., 2015) (implementation in (Evangelidis, 2013)), resulting in stabilized recordings without movement. For fura-2 fluorescence recordings, we first performed a single registration on the combined image (*f*_340_ + *f*_380_) and then applied the displacement vector field, computed by the registration algorithm, to both images (*f*_340_ and *f*_380_) separately. ROIs were defined manually at t_0_ and superimposed onto all subsequent images of the stabilized recording. At each time point t_*i*_, the fluorescence signal F was computed as the mean *f*_340_/*f*_380_ ratio of all pixels within a given ROI. When measuring Ca^2+^-dependent changes in GCaMP6f intensity, the fluorescence signal *F* was normalized with respect to a baseline before stimulation, computing the intensity change for the i^*th*^ time point as 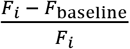. For clarity, linear baseline shifts were corrected in some example traces.

Seminiferous tubule contractions and transport of luminal content were visualized by reflected light microscopy of acute slices or whole-mount macroscopic tubule imaging, respectively. Data from both types of time-lapse recordings were analyzed and quantified as either flow strength or flow change (see below). For each frame at a given time point *t*^*i*^, the registration algorithm computed a flow or displacement vector field 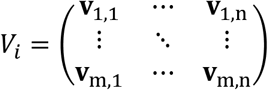, where **v**_1,1_ = (x, y) is a vector indicating strength and direction of the displacement of pixel (1,1) between time points *t*_0_ and *t*^*i*^. The average norm 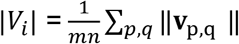 is a measure for the effort that is necessary to register the image at t_0_ to the image at *t*^*i*^. The flow field strength quantified by this measure is interpreted as the amount of visible changes that, dependent on the experiment, result from tubule contraction and / or luminal content movement. For analysis of contractions in acute seminiferous tubule slices (Figures 3, 4, 7), we quantified the *flow strength s*_*i*_ within an ROI as the average norm |*V*_*i*_| computed only for the **v**_p,q_ corresponding to pixels within the ROI defined at *t*_0_. For whole-mount macroscopic imaging of luminal content movement in intact tubule segments (Figure 5), we quantified the *flow change c*_*i*_ = *s*_*i*_ − *s*_*i*−1_ as the change of flow strength between two consecutive time points / frames. Here, *s*_*i*_ values were preprocessed by smoothing with a moving average filter. Results are reported as the AUC, i.e., the area under the *c*_*i*_ curve.

For analysis of *in vivo* data, we employed a custom set of ImageJ macros utilizing build-in functions of Fiji-ImageJ (Rueden et al., 2017; Schindelin et al., 2012). Widefield imaging data was first corrected for brightness fluctuation caused by a 50 Hz AC power supply. Here, we used the *bleach correction* plugin in histogram matching mode (Miura et al., 2014). Next, we applied Gaussian filter functions (*GausBlur* (5 px radius) and *Gaussian Blur3D* (x=0, y=0, z=5)). We calculated flow change via the *Gaussian Window MSE* function (sigma = 1; max distance = 3). Tubule selection used the polyline tool (line width adjusted to tubule diameter). Selected tubules ranged from 200 μm to 3.4 mm length. Next, flow fields of individual tubules were straightened. Average movement intensity was calculated from transversal line profiles (perpendicular to the straightened longitudinal axis of each tubule) and plotted as kymographs (space-time plots) to measure movement progression speed from linear regressions.

Multiphoton time-lapse imaging data was recorded in dual-channel mode, with (***i***) a target channel recording GCaMP6f fluorescence and some background signal (525\50 nm), and (***ii***) a background channel mainly recording autofluorescence (585\40 nm), allowing for background correction of the GCaMP6f signal using a dye separation routine. Slow constant movement in both channels was registered and removed to correct for steady drift. After Gaussian filtering (*GausBlur* (5 px radius); *Gaussian Blur3D* (x=0, y=0, z=5)), flow fields were calculated from the background signal. Again, flow change was calculated via the *Gaussian Window MSE* function (sigma = 1; max distance = 3).

## Additional information

### Competing interests

The authors declare no competing financial interests.

### Funding

This work was funded by the Deutsche Forschungsgemeinschaft (DFG, German Research Foundation) – 368482240/GRK2416 (NM & MSp); 412888997 (DF); 245169951 (AMa & MSp) – and by the Volkswagen Foundation (MSp, I/83533); MSp is a Lichtenberg Professor of the Volkswagen Foundation.

### Author contributions

D.F. and L.K. contributed equally and have equal right to list themselves first in bibliographic documents. D.F., L.K., A.Ma., J.S., and M.Sp. contributed substantially to the conception of this work. D.F., L.K., N.M., N.U., J.S., and M.Sp. designed experiments. D.F., L.K., N.M., F.B., A.Mi., R.M., N.U., M.St., and J.S. contributed to data acquisition and analysis. D.F., L.K., N.M., M.St., F.B., A.Ma., J.S., and M.Sp. contributed to data interpretation. M.St. and D.M. created new software used for contraction analysis. All authors have drafted the work and / or substantively revised it. All authors have approved the submitted version of the manuscript.

### Ethics

Mice were maintained and sacrificed according to European Union legislation (Directive 2010/63/EU) and recommendations by the Federation of European Laboratory Animal Science Associations (FELASA). All experimental procedures were approved by the State Agency for Nature, Environment and Consumer Protection (LANUV; protocol number / AZ 84-02.04.2016.A371).

## Acknowledgements

We thank Corinna Engelhardt, Jessica von Bongartz, and Stefanie Kurth (RWTH Aachen University) for assistance, Andreas Meinhardt and Jörg Klug (Justus-Liebig-University Giessen) for kindly providing detailed information about the mouse TPC cell culture protocol, Andrea Mietens and Ralf Middendorff (Justus-Liebig-University Giessen) for comments and suggestions, J. Ullrich Schwarzer (Andrology-Center, Munich) and Frank-Michael Köhn (Andrologicum, Munich) for providing human samples, and all members of the Spehr laboratory for discussions.

## Data and materials availability

All data is available in the main text or the supplementary materials. Previously unpublished source code for data analysis (quantification of tubular contractions, flow strength/change, Ca^2+^ signals) is available at: https://github.com/rwth-lfb/Fleck_Kenzler_et_al

---

**Figure 1 – figure supplement 1:**
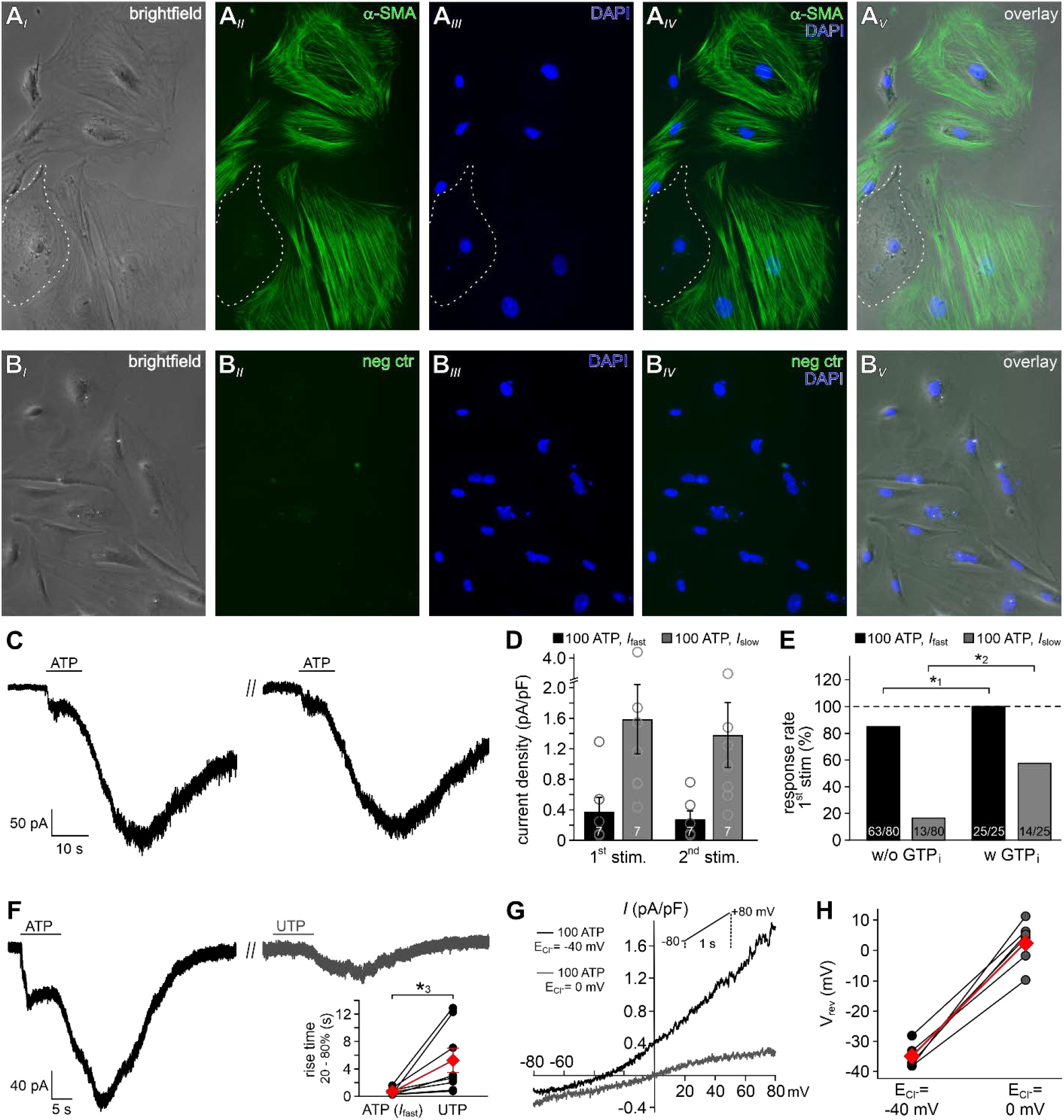
Mouse TPCs in primary culture display two distinct currents in response to extracellular ATP. (**A**) Representative phase contrast (**A**_*I*_) and epi-fluorescence images of TPCs *in vitro*. The vast majority of cultured cells are immunopositive for the TPC marker α-smooth muscle actin (α-SMA; green (**A**_*II*_). The dotted white line delimits one of the few α-SMA-negative cells. Cell count was based on nuclear staining (DAPI, blue (**A**_*III*_). Merged images (**A**_*IV-V*_) allow categorization. (**B**) Corresponding control images taken after the primary α-SMA antibody was omitted. (**C**) Representative whole-cell voltage-clamp recordings (V_hold_ = −80 mV) of ATP-induced inward currents in cultured mouse TPCs. Two components – a fast relatively small current and a delayed lasting current – are triggered repeatedly by successive ATP exposure (100 μM; 90 s inter-stimulus interval). Notably, we never observed a delayed slow current without a fast response. (**D**) Bar chart quantifying peak densities (mean ± SEM, circles show individual values) of the fast (*I*_fast_; black) and the delayed (*I*_slow_; grey) ATP-induced current components (1^st^ stimulation: *Ifast* 0.37 ± 0.2 pA/pF; *Islow* 1.58 ± 0.5 pA/pF; 2^nd^ stimulation *Ifast* 0.27 ± 0.1 pA/pF; *Islow* 1.37 ± 0.4 pA/pF). (**E**) Bar graph illustrating the frequency of *I*_fast_ (black) and *I*_slow_ (grey) occurrence upon ATP (100 μM) stimulation in absence (w/o) and presence (w) of GTP (500 μM) in the pipette solution, respectively. Asterisks denote statistically significant differences (*^1^*p* = 0.008, *^2^*p* = 0.0003; Fisher‘s exact test); n as indicated in bars. (**F**) Representative whole-cell voltage-clamp recordings (V_hold_ = −80 mV) of inward currents induced by ATP (100 μM) and UTP (100 μM), respectively. Whenever ATP triggers both *I*_fast_ and *I*_slow_ (left), *I*_slow_ is also induced by UTP (right). UTP-dependent currents develop significantly slower than ATP-evoked *I*_fast_ (inset; *^3^*p* = 0.03; paired *t*-test). (**G**, **H**) When measured during peak *I*_slow_, short (200 ms) voltage ramp (−80 to 80 mV) recordings reveal current-voltage relationships of ATP-induced currents at different Cl^−^ equilibrium potential (E_Cl_; −40 mV (**S**_**1**_/ **S**_**7**_; black trace) or 0 mV (**S**_**5**_/ **S**_**7**_; grey trace) (**G**). Note the corresponding shift in reversal potential (V_rev_) that is quantified in (**H**) (n = 6). Black / grey dots represent measurements from individual cells, red diamonds depict mean values.

**Figure 4 – figure supplement 1:**
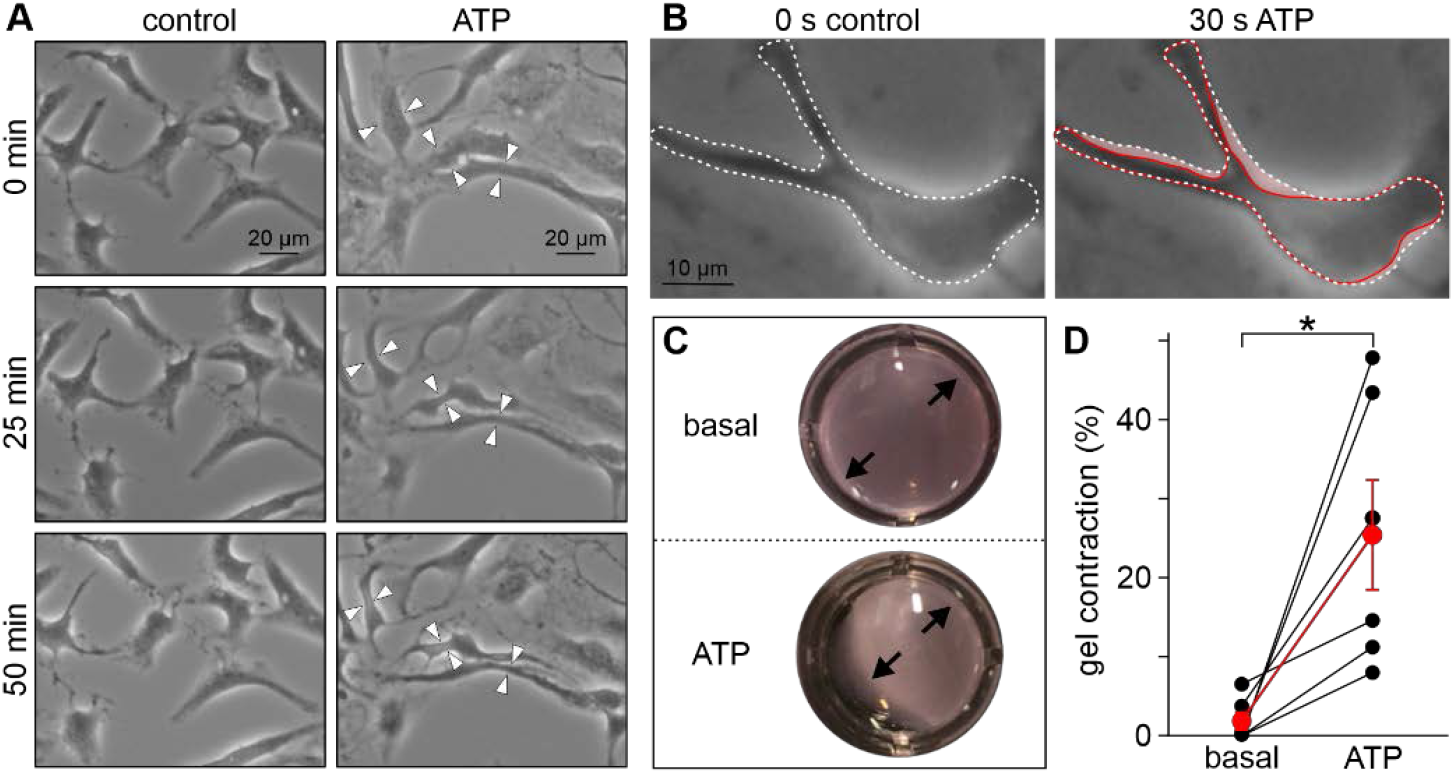
Purinergic stimulation mediates contractions in cultured human TPCs. (**A**) Phase contrast micrographs of human TPC in primary culture (Walenta et al., 2018) that were monitored (50 min) under either control conditions (left) or during treatment with ATP (right; 1 mM). White arrow heads denote regions where a substantial reduction in cell surface area upon ATP exposure becomes readily apparent. (**B**) Some TPCs already ‘shrink’ within 30 s of treatment. Cell contour is indicated before (dashed white line) and during (solid red line) ATP exposure (red ‘shadow’ illustrates the putative contraction). (**C**, **D**) 24 h collagen gel contraction assays (Ailenberg et al., 1990; Tung and Fritz, 1987) allow quantification of human TPC contractility *in vitro*. As exemplified in (**C**) and quantified in (**D**) ATP (1 mM) incubation of human TPCs that are embedded in collagen lattices mediates a massive reduction in gel area (red bar; 25.5 ± 7.0%, mean ± SEM; human TPCs from n = 3 patients, measured in duplicates). By contrast, gel size remains essentially unchanged under control conditions (black bar). Asterisk denotes statistical significance (*p* = 0.003; unpaired *t*-test).

**Figure 7 – figure supplement 1:**
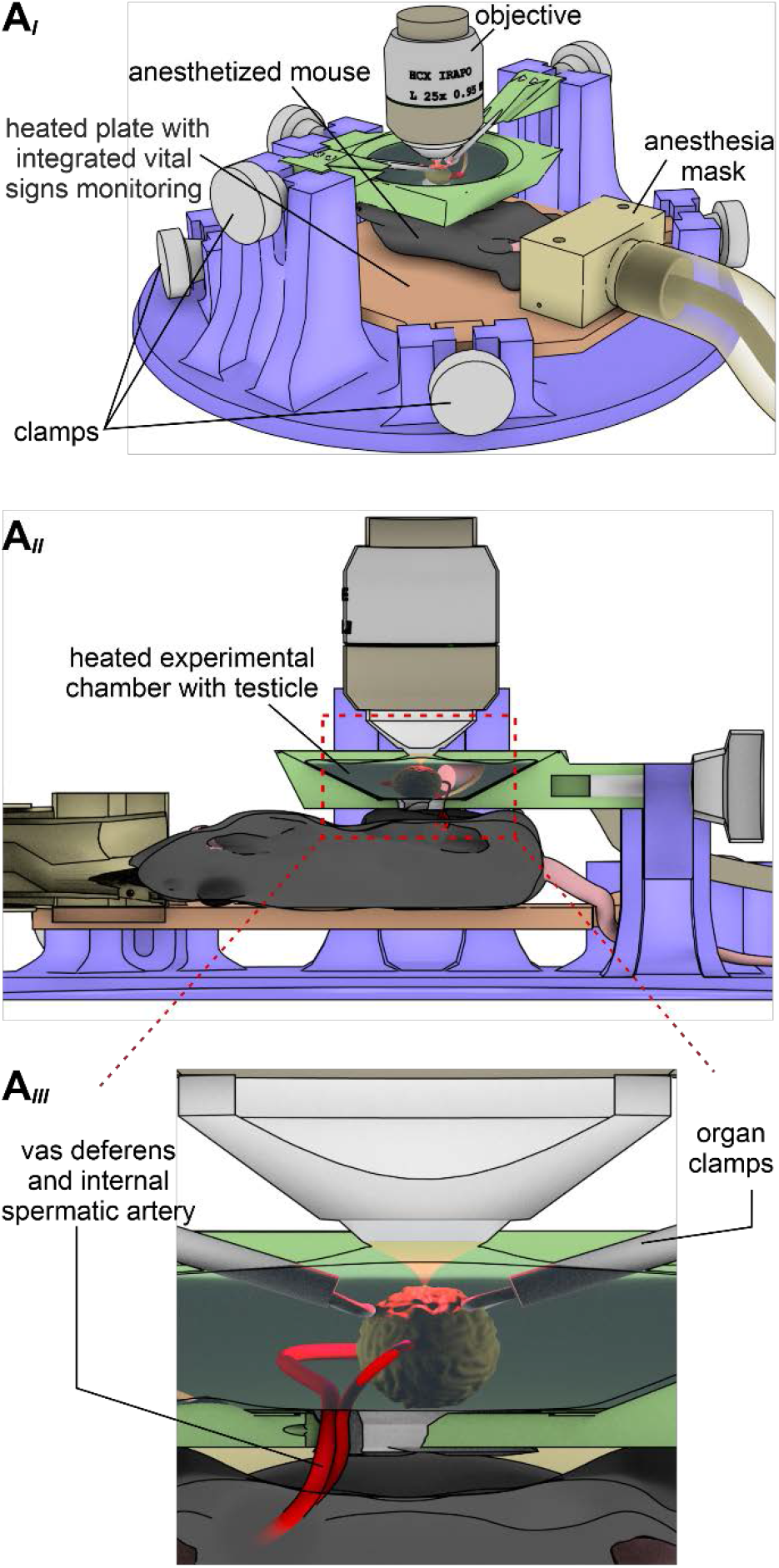
A custom-built 3D printed microscope stage enables simultaneous *in vivo* multiphoton imaging of Ca^2+^ signals and contractions in mouse seminiferous tubules. (**A**) Schematic drawing that illustrates the custom-built intravital imaging stage designed for *in vivo* recordings testicular activity. Three different views depict design details from different perspectives. A top view of the apparatus (**A**_*I*_) shows that several adjustable clamps allow exact positioning of the anaesthetized animal on a heated plate equipped for online vital sign monitoring. After centring one testicle in a heated (35°C) and saline-filled recording chamber (**A**_*II*_) within the microscope’s optical axis, a large working distance (~3 mm) infrared-optimized water-immersion objective (25x; 0.95 NA) enables multiphoton deep tissue imaging. For clarity, additional tubing that allows rapid exchange of oxygenated solution during experiments has been omitted. A close-up cartoon of the recording chamber (**A**_*III*_) illustrates that two micromanipulator-based moveable organ clamps enable precise (re)positioning of the testis as well as effective movement cancellation. Note that the *vas deferens* and internal spermatic arteries are kept intact to assure blood supply and fluid transport.

**Figure 7 – figure supplement 2:**
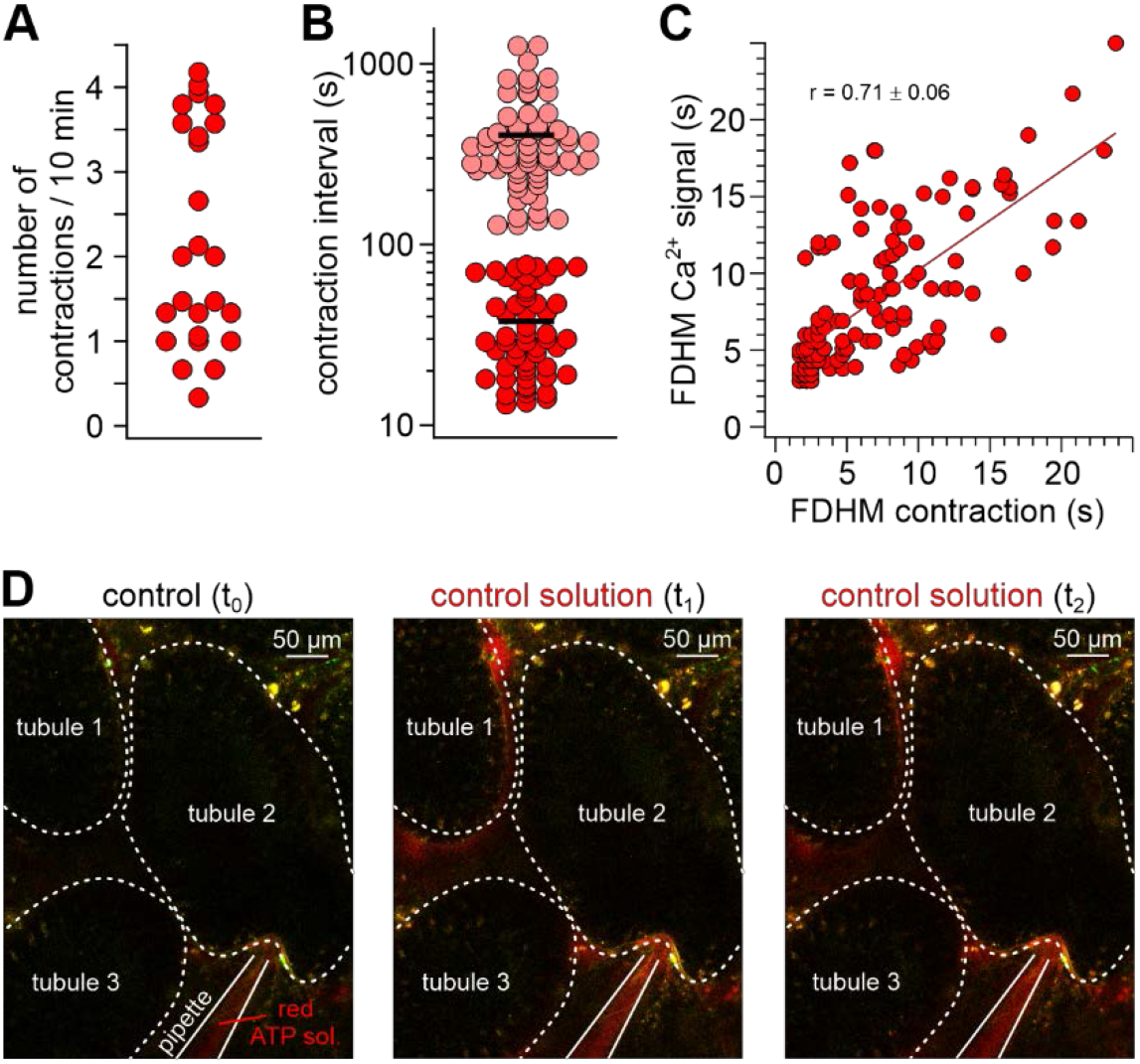
*In vivo* imaging of tubular activity in SMMHC-CreER^T2^ x Ai95D mice. (**A**–**C**) Quantitative analysis of spontaneous seminiferous tubule contractions *in vivo*. Dot plots depict (**A**) the number of contractions observed in a given tubule segment during 10 min windows of observation, and (**B**) the intervals between two consecutive contractions. Note that two distinct types of either relatively short (<80 s; red; 38 ± 21 s; mean ± SD) and long (>2 min; magenta; 403 ± 250 s; mean ± SD) intervals become apparent. Moreover, we never observed tubules that lacked contractions. (**C**) 2D dot plot shows positive correlation (Pearson correlation coefficient r = 0.71) between TPC Ca^2+^ signal durations (full duration at half-maximum; FDHM) and corresponding tubule contractions. (**D**) Example experiment that controls for pressure-dependent signal artifacts when seminiferous tubules are exposed to ‘puffs’ of solution. Three fluorescence images show outlines of three seminiferous tubule segments (white dotted lines) at different time points (t_0_ – t_2_). Aside the addition of a red fluorescent dye to the solution in the pipette, compositions of bath and pipette solution match. Note that puffs of stained solution (t_1_ & t_2_) cause neither contractions, nor Ca^2+^ signals.

**Movie S1: SMMHC-CreER^T2^ mice allow inducible TPC-specific expression of genetically encoded fluorescent reporter proteins.** After tamoxifen injections, SMMHC-CreER^T2^ x Ai14D male offspring express tdTomato (red) in both TPCs and vascular smooth muscle cells. Movie shows the 3D reconstruction of an intact and cleared (CLARITY (Chung and Deisseroth, 2013)) 6 × 3 × 1.5 mm testis sample with nuclei labelled by DRAQ5 (blue).

**Movie S2: Quasi-simultaneous recording of peritubular Ca^2+^ signals and seminiferous tubule movement.** A representative seminiferous tubule section (250 μm) is stimulated with ATP (100 μM, 10 s). After fura-2 bulk loading, ratiometric fluorescence imaging (*f*_340_/*f*_380_) reveals relative changes in Ca^2+^ concentration (rainbow colour map; blue, low Ca^2+^; red, high Ca^2+^) in a peripheral band of putative TPCs at the tubule’s edge. Since each image acquisition cycle (1 Hz) captures two fluorescence (Exλ340; Exλ380) and one reflective light image (brightfield), time-lapse recordings allow parallel physiological phenotyping of both seminiferous tubule Ca^2+^ responses and movement (shown sequentially for clarity).

**Movie S3: Both ATP-induced seminiferous tubule Ca^2+^ responses and contractions are dose-dependent.** A representative seminiferous tubule section (250 μm; fura-2 bulk loading) is stimulated with increasing ATP concentrations (1–1000 μM, 10 s). Ratiometric fluorescence imaging (*f*_340_/*f*_380_) reveals relative changes in Ca^2+^ concentration (rainbow colour map; blue, low Ca^2+^; red, high Ca^2+^) in putative TPCs. Quasi-simultaneous time-lapse recording of fluorescence (Exλ_340_; Exλ_380_) and brightfield (reflective light) images illustrates that both seminiferous tubule Ca^2+^ signals and contractions (shown sequentially for clarity) are dose-dependent and share an ATP threshold concentration of approximately 1 μM.

**Movie S4: ATP stimulation triggers movement of luminal content in intact seminiferous tubules.** Brightfield time-lapse recording of an intact isolated seminiferous tubule (field of view shows cycle stages II and III) challenged by brief focal ATP perfusion (100 μM, 10 s). The spatial extent of the stimulation zone had been defined by prior perfusion with a dye solution (food color).

**Movie S5: ATP stimulation triggers transient Ca^2+^ signals in TPCs of intact seminiferous tubules.** Fluorescence time-lapse recording of an intact seminiferous tubule (field of view shows cycle stage II) isolated from a mouse selectively expressing GCaMP6f in TPS (SMMHC-CreER^T2^ x Ai95D male offspring). Fluorescence imaging (ΔF/F) during brief focal ATP perfusion (100 μM, 10 s) – the spatial extent of the stimulation zone had been defined by prior perfusion with a dye solution (food color) – reveals relative changes in TPC Ca^2+^ concentration (rainbow colour map; blue, low Ca^2+^; red, high Ca^2+^).

**Movie S6: *In vivo* multiphoton microscopy demonstrates spontaneous Ca^2+^ signals in mouse TPCs.** Spontaneous seminiferous tubule *in vivo* activity monitored in SMMHC-CreER^T2^ x Ai95D mice. Intravital multiphoton fluorescence time-lapse imaging (ΔF/F, 2 Hz) reveals coordinated changes in TPC Ca^2+^ concentration (rainbow colour map; blue, low Ca^2+^; red, high Ca^2+^) among one of three seminiferous tubules in the field of view (591 μm × 591 μm).

**Movie S7: Coordinated contractile movements ensure luminal sperm transport *in vivo*.** Intravital *en-face* brightfield imaging illustrates spontaneous contractions and luminal movement in seminiferous tubules of adult mice. Low (1.5 mm × 1.4 mm field of view) and high-magnification time-lapse recordings reveal that contractions and luminal content propulsion are routinely observed *in vivo*. Note the unobstructed blood flow within testicular vessels.

**Movie S8: Focal ATP stimulation triggers peritubular Ca^2+^ signals and seminiferous tubule contractions *in vivo*.** Intravital multiphoton fluorescence time-lapse imaging in SMMHC-CreER^T2^ x Ai95D mice. Overlay of two detection channels (ΔF/F, GCaMP6f, green; Alexa Fluor 555, red). Stimulus solution (containing Alexa Fluor 555 (4 μM) and ATP (1 mM)) is puffed from a glass micropipette, which penetrated the *tunica albuginea* to target the interstitial space. Changes in TPC Ca^2+^ concentration are color-coded (black, low Ca^2+^; green, high Ca^2+^). Note that typically such contractions / Ca^2+^ signals do not occur when ATP is omitted from the ‘puff’ solution (data not shown).

